# Genome-wide 5-hydroxymethylcytosine (5hmC) emerges at early stage of *in vitro* hepatocyte differentiation

**DOI:** 10.1101/629493

**Authors:** Jesús Rafael Rodríguez-Aguilera, Szilvia Ecsedi, Marie-Pierre Cros, Chloe Goldsmith, Mariana Domínguez-López, Nuria Guerrero-Celis, Rebeca Pérez-Cabeza de Vaca, Isabelle Chemin, Félix Recillas-Targa, Victoria Chagoya de Sánchez, Héctor Hernández-Vargas

## Abstract

How cells reach different fates despite using the same DNA template, is a basic question linked to differential patterns of gene expression. Since 5-hydroxymethylcytosine (5hmC) emerged as an intermediate metabolite in active DNA demethylation, there have been increasing efforts to elucidate its function as a stable modification of the genome, including a role in establishing such tissue-specific patterns of expression. Recently we described TET1-mediated enrichment of 5hmC on the promoter region of the master regulator of hepatocyte identity, HNF4A, which precedes differentiation of liver adult progenitor cells *in vitro*. Here we asked whether 5hmC is involved in hepatocyte differentiation. We found a genome-wide increase of 5hmC as well as a reduction of 5-methylcytosine at early hepatocyte differentiation, a time when the liver transcript program is already established. Furthermore, we suggest that modifying s-adenosylmethionine (SAM) levels through an adenosine derivative could decrease 5hmC enrichment, triggering an impaired acquisition of hepatic identity markers. These results suggest that 5hmC is a regulator of differentiation as well as an imprint related with cell identity. Furthermore, 5hmC modulation could be a useful biomarker in conditions associated with cell de-differentiation such as liver malignancies.

**Graphical Abstract:** It has been suggested that 5-hydroxymethylcytosine (5hmC) is an imprint of cell identity. Here we show that commitment to a hepatocyte transcriptional program is characterized by a demethylation process and emergence of 5hmC at multiple genomic locations. Cells exposed to an adenosine derivative during differentiation did not reach such 5hmC levels, and this was associated with a lower expression of hepatocyte-markers. These results suggest that 5hmC enrichment is an important step on the road to hepatocyte cell fate.

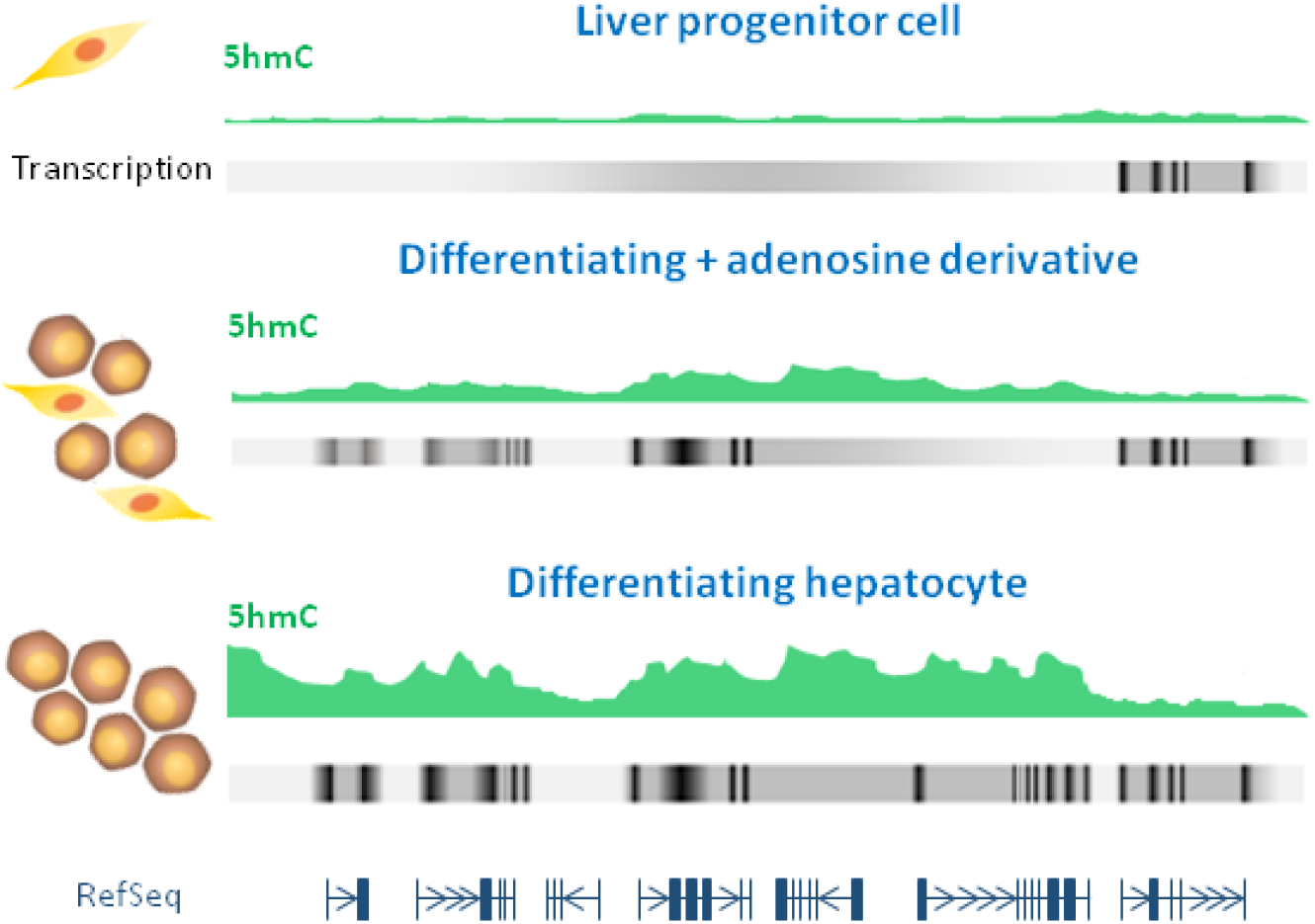

## Introduction

Chronic liver disease comprises multiple different pathologies, such as non-alcoholic fatty liver disease, liver fibrosis and cirrhosis. Commonly these pathologies are the background to develop liver cancer [Sia et al., 2017], which encompass a heterogeneous group of malignant tumours with different histological features, and unfavourable prognosis. Hepatocellular carcinoma (HCC) and intrahepatic cholangiocarcinoma (iCCA) are the most common primary cancers of the liver [Ervik et al., 2016], which was estimated as the seventh in incidence and fourth highest cause of cancer-related death worldwide in 2018 [The, 2018].

Several studies point to adult hepatocytes as the cell of origin of liver cancer [Mu et al., 2015; Shin et al., 2016], through de-differentiation into hepatocyte precursor cells that later become HCC cells or through *trans*-differentiation into biliary-like cells, which give rise to iCCA [Sia et al., 2017]. In spite of their different aetiologies, chronic liver damage is a shared factor in HCC and iCCA development. This constant insult results in high cell turnover, and aberrant genetic and regulatory landscapes that favours liver cell transformation [Castelli et al., 2017]. As transformation takes place, progenitor-like cells accumulate, in a process known as ductular reaction [Mu et al., 2015] where cholangiocytes could also *trans*-differentiate into stress-resistant hepatocytes to help in liver repopulation of damaged regions [Manco et al., 2019]. This underscores the importance of understanding hepatocyte differentiation as a key process in chronic liver injury, where cell identity is lost.

With the advance on the understanding of chromatin organization of the eukaryotic genome it has been clear that epigenetics influences both, normal human biology and disease [Recillas-Targa, 2014], being DNA methylation one of the most studied epigenetic modifications. DNA methyl-transferases (DNMTs) establish 5-methylcytosine (5mC) from S-adenosylmethionine (SAM), the principal methylating agent in the body derived from the methionine cycle [Tehlivets et al., 2013]. In humans, the liver is the organ with the highest turnover of SAM and metabolism of methionine [Mato et al., 2002]. It has been suggested, that adenosine could be able to modulate SAM methylation in the liver, promoting either the metabolic flow of methyl group transfer-reactions or their inhibition by inducing S-adenosylhomocysteine (SAH) accumulation [Chagoya de Sanchez et al., 1991].

As opposed to DNMTs, Tet Methylcytosine Dioxygenases (TET) are involved in the oxidation of methylated cytosine in DNA forming 5-hydroxymethylcytosine (5hmC) and together, with the base excision DNA repair machinery, leading to active cytosine demethylation [Kriaucionis and Heintz, 2009; Tahiliani et al., 2009]. Moreover, 5hmC is emerging as a stable modified base with unique functions that suggests that the dynamic distribution of 5hmC could be an acquired “imprint” of cell identity during adult progenitor cell differentiation in several tissues (reviewed in [Ecsedi et al., 2018]) including liver. Indeed, using an *in vitro* model of hepatocyte differentiation, we recently described a switch in 5mC/5hmC state of promoter 1 (P1) of the master transcription factor (TF) of hepatocyte identity *HNF4A* [Ancey et al., 2017].

The capacity to modulate epigenetic modifications, offers an opportunity to develop new strategies for the early prevention and treatment of diseases [Li et al., 2018]. An adenosine derivative, IFC-305 (UNAM Patent 207422), administered in a cirrhotic rat model has previously been able to recover SAM levels abrogated in cirrhosis which triggers the re-establishment of 5mC and 5hmC in this condition [Rodriguez-Aguilera et al., 2018]. IFC-305 also has several hepatoprotective properties (reviewed in [Rodríguez-Aguilera et al., 2019]), such as, reduction of fibrosis and amelioration of liver function through regulation of transcriptome [Perez-Carreon et al., 2010], promotion of cell cycle recovery in the liver [Chagoya de Sanchez et al., 2012], anti-inflammatory effects during cirrhosis [Perez-Cabeza de Vaca et al., 2018], prevention of hepatic stellate cells (HSCs) activation [Velasco-Loyden et al., 2010], and anti-carcinogenic effects [Velasco-Loyden et al., 2017].

Here, we asked whether 5hmC is present and/or redistributed in the genome of differentiating hepatocytes. We describe 5hmC genomic enrichment and its relationship with gene expression. Moreover, we show that 5hmC accumulation and hepatocyte differentiation are impaired by perturbing SAM metabolism using IFC-305.

## Results

### HepaRG cells differentiated for 1 week express hepatocyte markers

HepaRG cells are bipotent liver progenitor cells that differentiate *in vitro* after 4 weeks into either hepatocytes or cholangiocytes. Our group previously found a TET1-dependent switch from methylated to hydroxymetylated DNA status at *HNF4A* promoter P1 in HepaRG cells, triggering differentiation at one week of cell culture [Ancey et al., 2017]. In order to determine the gene expression profile at this stage of hepatocyte differentiation (Figure 1A), RNA was isolated and a transcriptome analysis was performed to identify differentially expressed genes (DEGs) (Figure 1B). We found 4175 DEGs upon one week of differentiation. While down-regulated genes (n=2066 probes, corresponding to 1772 hg19-annotated genes) were related to lymphoblasts and endothelial cells (Figure 1C), over-expressed genes (2109 probes, corresponding to 1822 hg19-annotated genes) were highly associated with liver and foetal liver cells (Figure 1D). In addition, we found that over-expressed genes were enriched in targets of the *HNF4A* transcription program (Supplementary Figure S1A). Pathways most related with these genes included biological oxidation and metabolism related signalling pathways (Supplementary Figure S1B). Gene ontologies revealed fatty acids, regulation of lipids, and triglyceride homeostasis, metabolic processes as well as oxidoreductase, and endopeptidase and alcohol dehydrogenase activities (Supplementary Figures S1C and S1D). On the other hand, down-regulated genes in differentiating cells are associated with *E2F4* transcriptional program (Supplementary Figure S1E), signalling pathways involved in cell cycle progression, biological process related with DNA metabolism and replication, and molecular functions implicated in DNA dependent ATPase activity (Supplementary Figures S1F - S1H). We assessed expression levels of hepatocyte markers over-expressed in transcriptome data and validated the overexpression of *HNF4A* P1 isoforms, *GSTA*, and *ALDOB* (Figures 1E-1H).

**Figure 1.**
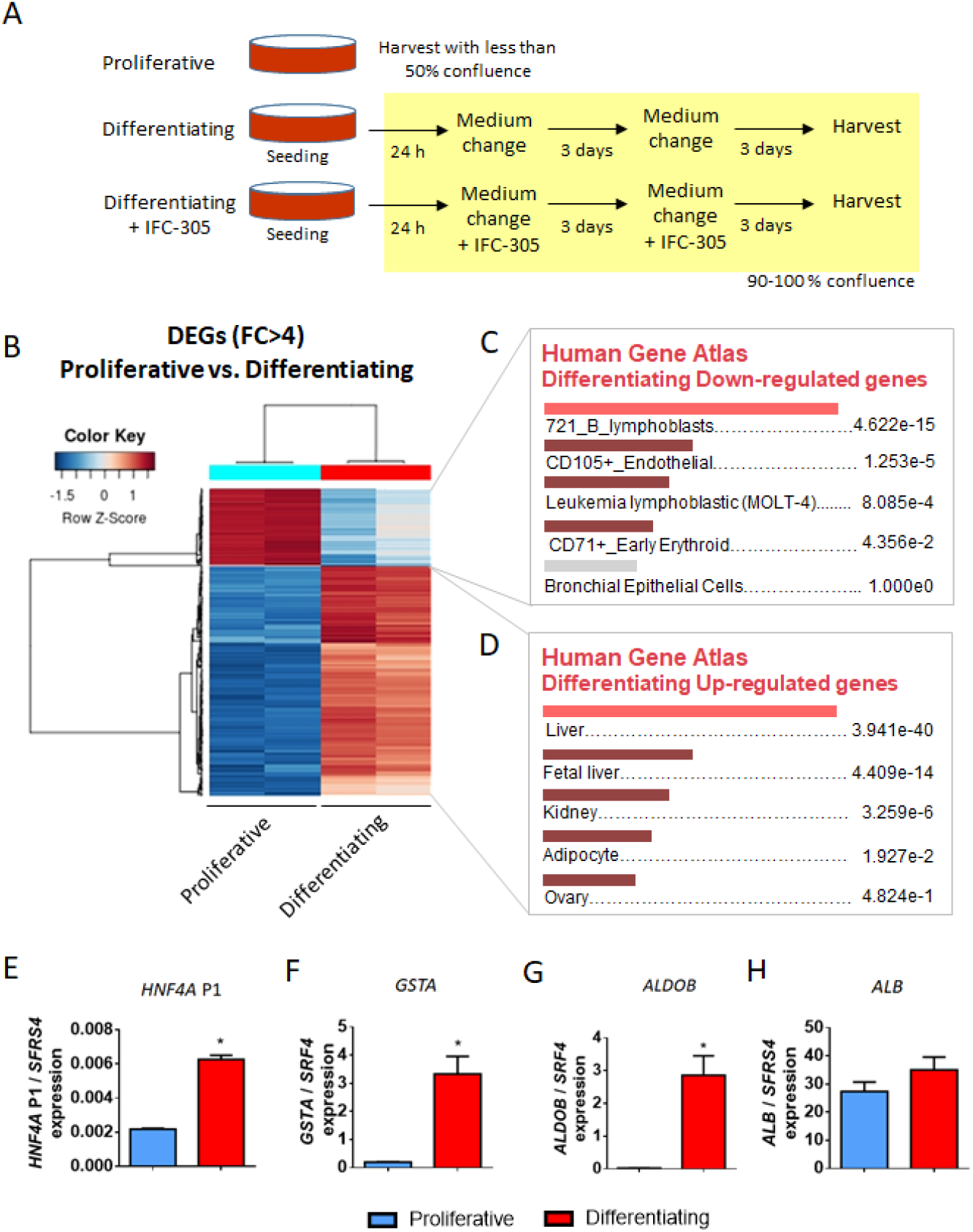
Liver transcription program is expressed in HepaRG cells at 1 week of differentiation. (A) HepaRG differentiation model. For proliferative (progenitor) condition, cells were seeded and trypsinized after reaching 50% confluence; for differentiating conditions, cells were seeded at 70-80% confluence in order to reach 100% confluence 24 h after seeding. (B) Transcriptome was analyzed in both conditions. Heatmap represents differentially expressed genes (DEGs) with fold change greater than four. Cell/tissues types associated with genes down-regulated (C) and up-regulated (D) in differentiating cells (EnrichR), adjusted p-values are shown. (E-H) Expression of hepatocyte markers was validated by RT-qPCR, data represent mean ± SEM 3 independent cultures/condition; *Statistical difference (p < 0.05).

Altogether, these results indicate that after 1 week of differentiation, HepaRG cells have turned on a hepatocyte-like expression program, while proliferative related genes become progressively silenced.

### HepaRG differentiation involves an early 5hmC landscape remodelling

Considering that at 1 week of HepaRG differentiation there is a TET1-mediated 5hmC enrichment on *HNF4A* promoter P1 [Ancey et al., 2017] and the transcriptome already reflects a hepatocyte-like profile, we chose this time point to assess 5hmC levels of the HepaRG cell line compared to its proliferative state. Immunostaining against 5hmC reveals the presence of this modified cytosine in differentiating cells, in contrast with its almost complete absence in proliferative HepaRG cells (Figure 2A and Supplementary Figure S2).

**Figure 2.**
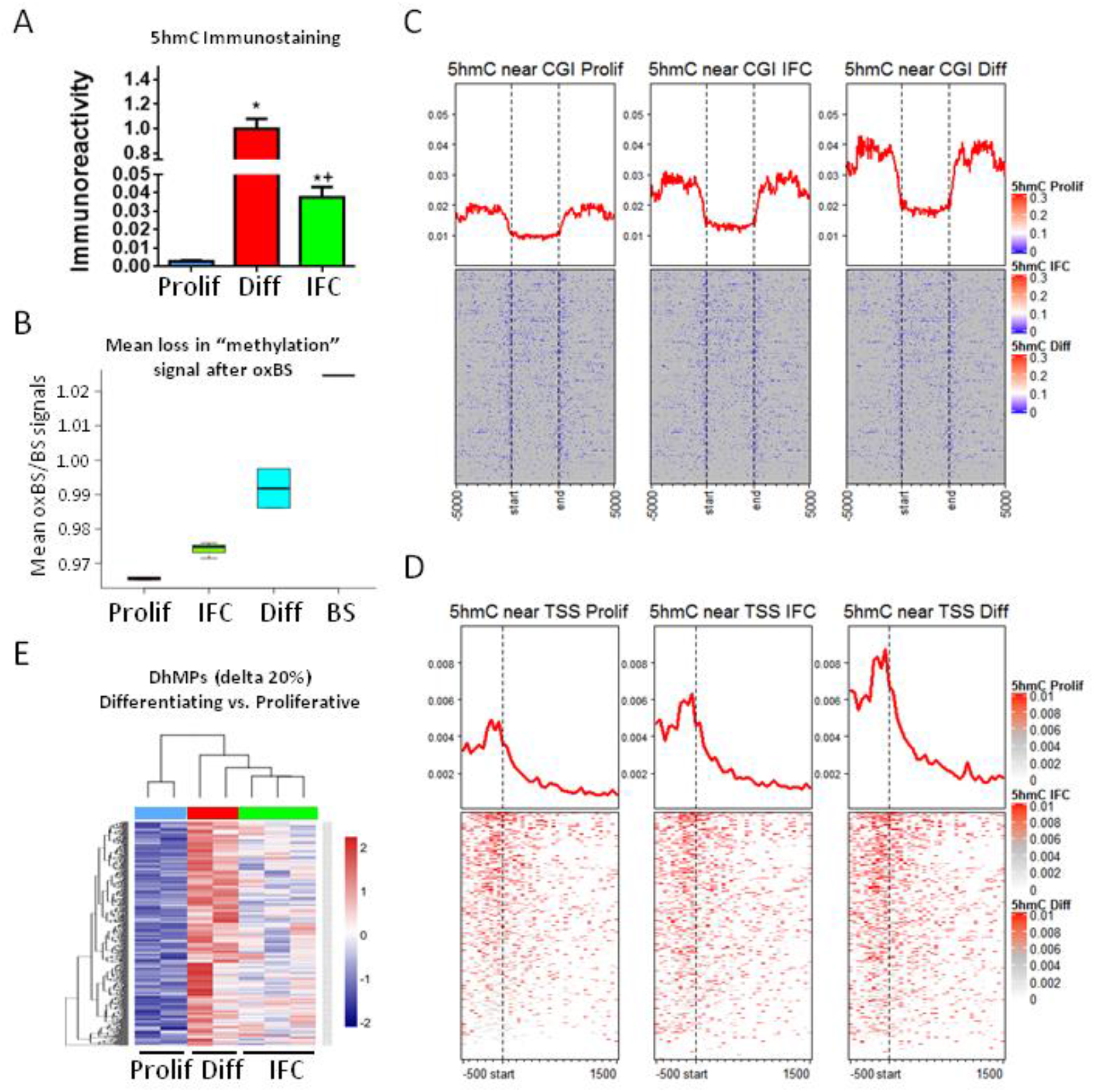
5hmC genome-wide enrichment during hepatocyte differentiation. (A) Immunofluorescence of 5hmC in proliferative (Prolif), differentiating (Diff) and differentiating + IFC-305 (IFC) HepaRG cells. Quantification barplots of 5hmC signal are based on mean ± SEM from 3 fields/group. *Statistical difference (p < 0.05), or *^+^ (p < 0.05) when compared with proliferative and differentiating cells. (B) The same conditions were assessed for base-resolution 5mC/5hmC, using BS and OxBS followed by EPIC beadarray hybridization (see Methods). Proportion of signal loss after oxidation (mean oxBS signal / mean BS signal) was used as an estimation of global 5hmC in each condition. BS represents the variation between two BS technical replicates. (C) Global distribution of 5hmC for one representative sample of each condition, according to CpG islands (CGI) (C) and transcription start sites (TSS) (D). In both cases, 5hmC levels are averaged across all hg19-annotated genomic regions. (E) Heatmap showing hydroxymethylome comparison between differentiating and proliferative cells. Differentially hydroxymethylated positions (DhMPs) were filtered by the magnitude of change in methylation (delta beta) of at least 20% and p-adjusted value < 0.05. Two independent cultures were used for proliferative and differentiating cells, and three independent cultures for differentiating + 1 mM IFC-305.

Next, we proceeded to evaluate the hydroxymethylome at base resolution. At this point it is worth mentioning that conventional DNA bisulfite conversion is not able to distinguish 5mC and 5hmC. An alternative to evaluate both modifications independently, is the use of oxidative bisulfite; which allows for the identification of 5mC through the oxidation of 5hmC to 5-formylcytosine (5fC) with KO_4_Ru [Booth et al., 2013]. Therefore, we isolated DNA from proliferative and differentiating HepaRG cells and performed oxidative and conventional bisulfite conversion (oxBS and BS, respectively), followed by hybridization on Infinium EPIC arrays.

Using only oxBS signal, we observed the expected distribution of 5mC in CpG islands (CGIs) and transcription start sites (TSS) (Supplementary Figure S3). Although no global differences were evident between conditions, we identified 3351 differential methylated positions (DMPs) displaying lower methylation after one week of differentiation (Supplementary Figure S4A).

Roughly, a loss of signal after oxBS relative to conventional BS indicates the presence of 5hmC. While analysis of genomic data showed almost no signal loss between oxBS and BS in proliferative cells, a significant global loss was evident in differentiating cells suggesting a global gain in 5hmC during the first week of differentiation (Figure 2B). Such gain in 5hmC was consistent, regardless of relative location across CGIs and TSS (Figure 2C and 2D).

Next, we studied 5hmC by directly comparing oxBS and BS data. While no significant global signal in 5hmC was observed in proliferative cells (Figure 2B), we identified a gain of 5hmC at 11766 sites in differentiating cells, defined by a significant reduction of methylation signal after oxBS of at least 10% (5hmC peaks). Differential hydroxymethylation performed at these sites revealed 6952 differentially hydroxymethylated positions (DhMPs) between proliferative and differentiating cells (Figure 2E). In addition, we identified 2482 differentially hydroxymethylated regions (DhMRs). Among these, a 3-CpGs region on *HNF4a* was identified as well as a 21-CpGs region, a 5-CpGs region and two 3-CpGs regions on *IDH3G, TET1* and *TET3* genes, respectively (Supplementary Figures S5 – S9). All of these genes are involved themselves in establishment of 5hmC.

Together, these results show that while 5hmC is poorly present at the HepaRG progenitor stage, it is enriched at multiple genomic locations upon entering hepatocyte differentiation.

### Genomic context of differentially hydroxymethylated sites associated with early hepatocyte differentiation

In order to know whether there is a relationship between 5hmC enrichment and gene expression, we compared the nearest associated gene of DhMPs with DEGs associated with one week of differentiation (described above). Out of 6952 DhMP-associated genes, 522 and 482 fall near an up- or a down-regulated DEG, respectively (Figure 3A). Globally, however, there was no difference in the genomic 5hmC distribution in DEGs (up- or down-regulated) compared to control genes (Figure 3B).

**Figure 3.**
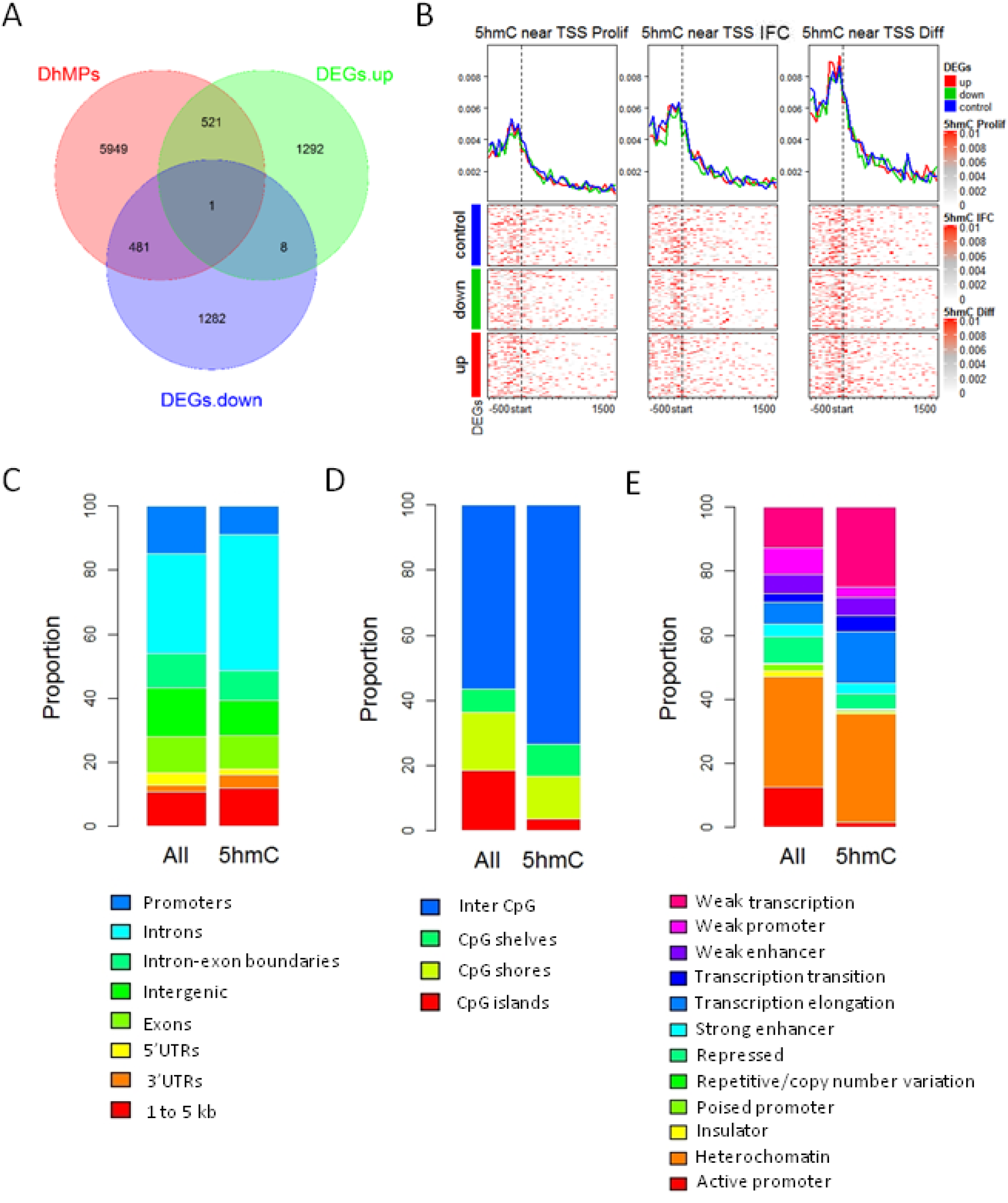
Association between 5hmC and differential expression. (A) Venn diagram representation of the overlaps between 5hmC differentiated positions (DhMPs) and differential expressed genes (DEGs). For this analysis, DhMPs were annotated to its nearest gene. (B) Global distribution of 5hmC according to transcription start sites. 5hmC levels are averaged across all hg19-annotated genomic regions, in turn divided into control, up-regulated, or down-regulated genes. DhMPs were also annotated according to gene features (C), distribution according to CpG islands (D), and HepG2 chromatin states (ChromHMM) (D). For each genomic context, distribution is shown separately for all DhMPs, and all EPIC beadarray probes, as a control. Two independent cultures were used for proliferative and differentiating cells, and three independent cultures for differentiating + 1 mM IFC-305.

DhMPs associated with increased gene expression were related with liver and fetal liver cell types, and enriched in pathways involved in androgen receptor (*AR*) and Nuclear Factor Erythroid 2-Related Factor 2 (*NFE2L2*) transcription programs as well as part of metabolism signalling pathways (Supplementary Figures S10A – S10C). DhMPs associated with down-regulated genes, were related with B lymphoblasts and enriched in pathways involved in cell cycle control, such as the *E2F4* transcription program (Supplementary Figures S10D - S10F).

Next, in order to explore a less evident role of 5hmC enrichment on gene expression, we compared distribution of DhMPs relative to different genomic annotations. DhMPs were enriched in intronic regions, and depleted from promoters and CGIs (Figure 3C and 3D). In addition, when we assessed genomic annotated features of liver HepG2 cells, we found an over-representation of DhMPs in weak transcription and transcription elongation loci (Figure 3E).

In summary, these results show that only 17% of 5hmC enrichment during HepaRG differentiation can be directly associated with gene expression changes at the neighbouring genomic location.

### Methyl-donor perturbation has repercussions for hepatocyte differentiation

Considering the genome-wide increase in 5hmC upon early HepaRG differentiation, we evaluated if a disruption in 5hmC levels could modify the hepatocyte differentiation process. To assess this question, we used a previously described adenosine derivative, IFC-305, which is able to ameliorate liver function and reduce collagen accumulation during cirrhosis. IFC-305 recovers both 5mC and 5hmC, abrogated during cirrhosis, improving SAM availability [Rodriguez-Aguilera et al., 2018].

Firstly, we evaluated HepaRG cell viability in response to IFC-305 exposure and we found that viability was not affected at concentrations of up to 1 mM for 1 week of differentiation (Supplementary Figure S11). With this maximum concentration, we analysed the expression of hepatocyte markers in response to different concentrations of IFC-305 and noticed 1 mM triggered lower values of *ALDOB* and *GSTA* in comparison with non-treated differentiating cells (Figure 4A). Albumin expression did not change, neither on differentiating non-treated cells nor after IFC-305 treatment, except for a reduction in expression levels with 5 mM IFC-305 (Figure 4A) compared with proliferative cells. Regarding *HNF4A* isoform regulated by P1 gene promoter, its expression showed the same increment level with 0.2 and 1 mM IFC-305 and non-treated differentiating cells, but a concentration of 5 mM generated an over-expression of this isoform compared with proliferative cells (Figure 4B). On the other hand, *HNF4A* isoforms regulated by P2 gene promoter increased its expression only with 0.2 and 1 mM IFC-305 on differentiating cells in comparison with proliferative cells (Figure 4B). However, at the protein level, 1 mM IFC-305 was associated with a reduced HNF4α nuclear immunoreactivity signal in differentiating cells compared with non-treated differentiating ones (Supplementary Figure S12).

**Figure 4.**
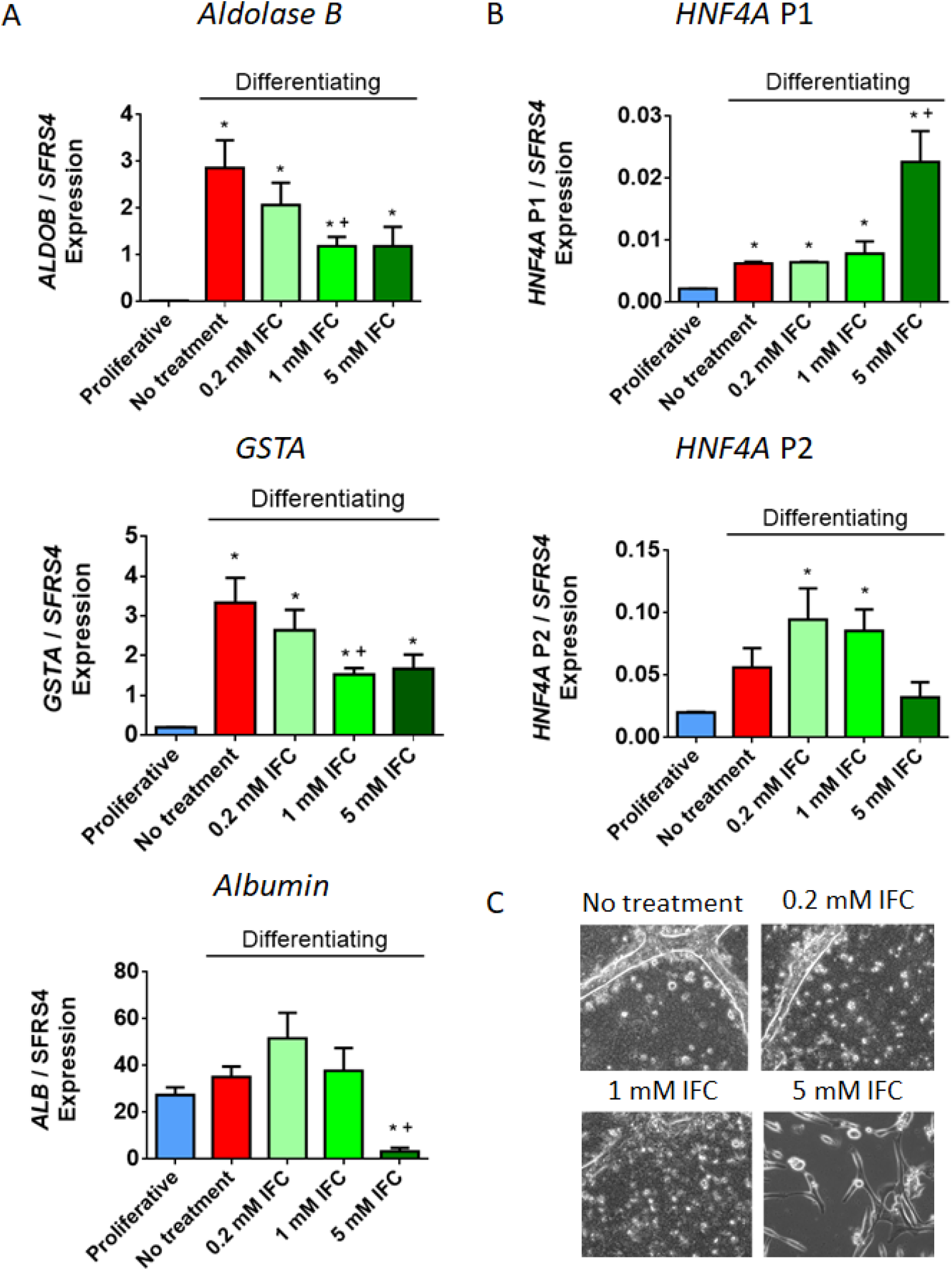
Impaired hepatocyte marker expression upon adenosine derivative exposure. (A) Expression of hepatocyte markers was assessed by RT-qPCR, data represent mean ± SEM 3 independent cultures/condition. (B) Expression of isoforms from HNF4A promoters 1 and 2 was assessed by RT-qPCR, data represent mean ± SEM 3 independent cultures/condition. (C) Phase contrast images showing HepaRG differentiated phenotype after 20 days of exposure to an increasing gradient of IFC-305. Representative 20x magnification images from seven independent cultures are shown. *Statistical difference (p < 0.05). ^*+^Statistical difference (p < 0.05) when compared with proliferative and differentiating non-treated cells.

This pattern of expression of hepatocyte markers suggests that HepaRG exposure to IFC-305 during differentiation influences hepatocyte maturation. Exploring this possibility, we extended the differentiation model to a longer period (20 days). At 13 days, differentiating non treated cells show hepatocyte-like cell morphology, emergence of small polygonal cells with increased granularity and organized in well-delineated trabeculae, separated by bright canaliculi-like structures (Supplementary Figure S13). IFC-305 delays this phenotype in a concentration-dependent manner, with 1 mM IFC-305 treated cells showing a phenotypic delay of at least one week, and 5 mM IFC-305 retaining a proliferative-like phenotype up to day 20 of differentiation (Figure 4C and Supplementary Figure S13). Such phenotypic variation, reduced levels of hepatocyte markers (i.e. *HNF4A* isoforms regulated by P1 gene promoter, *AldoB* and *GSTA*) and increased expression of *HNF4A* isoforms regulated by P2 gene promoter, suggest that IFC-305 is able to delay hepatocyte differentiation.

### Methyl-donor perturbation disrupts the 5hmC landscape associated with hepatocyte differentiation

Proliferative HepaRG cells were exposed to 1 mM IFC-305 in differentiating conditions during 1 week. Under these conditions, 5mC reduction and 5hmC accumulation are impaired, as assessed by IF and global oxBS and BS analyses (Figures 2A and 2B, #Supplementary Figure S4B). Similarly, only a fraction of differentiation-related DhMPs are detected in presence of IFC-305 using base-resolution methylation bead arrays (Figure 2E). Despite this attenuated phenotype, differential hydroxymethylation was identified at 8460 gene associated CpGs sites (Figure 5A), and 1890 regions, relative to proliferative cells. Comparing these genes with up-regulated DEGs in differentiating cells, we found an overlap of 173 genes which are highly associated with liver expression, according to the ARCHS4 Tissues data base (Supplementary Figure 14).

**Figure 5.**
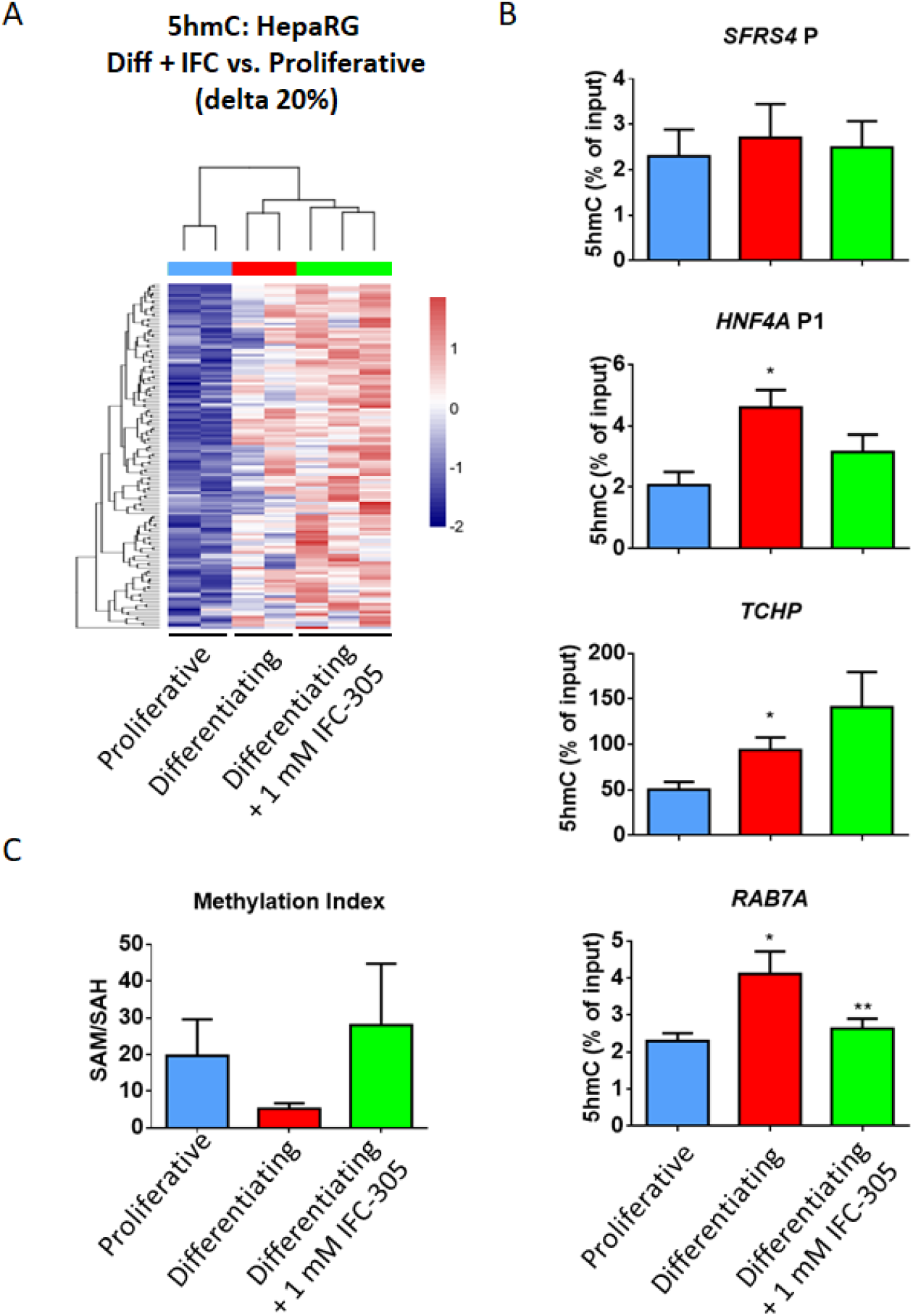
Reduced 5hmC enrichment under methyl-donor exposure. (A) 5hmC in proliferative, differentiating and differentiating + IFC-305 HepaRG cells, as assessed using BS and OxBS followed by EPIC beadarray hybridization (see methods and Figure 2). Heatmap showing hydroxymethylome comparison between differentiating + IFC-305 and proliferative cells. Differentially hydroxymethylated positions (DhMPs) were filtered by the magnitude of change in methylation (delta beta) of at least 20% and p-adjusted value < 0.05. Two independent cultures were used for proliferative and differentiating cells, and three independent cultures for differentiating + 1 mM IFC-305. (B) Validation of 5hmC changes. 5hmC enrichment was assessed by hMeDIP and qPCR; *SFRS4* gene promoter was used as non-5hmC enriched control region, HNF4A promoter P1 was used as 1 week differentiation 5hmC enrichment positive control, data represent mean ± SEM 3 independent cultures/condition; *Statistical difference (p < 0.05) compared with proliferative cells. **Statistical difference (p < 0.05) compared with differentiating cells. (C) Content of S-adenosylhomocysteine (SAH) and S-adenosylmethionine (SAM) was assessed using HPLC in different experimental conditions and generated by standards. Barplots indicate the methylation index, calculated as the SAM/SAH ratio, mean ± SEM from 4 cultures/condition.

In order to validate the increment of 5hmC through the differentiation process, we performed hydroxymethylated DNA immunoprecipitation assays (hMeDIP) on selected regions with the highest 5hmC increase. We assessed two different regions in the vicinity of *TCHP and RAB7A*. Furthermore, we evaluated the region where a switch in 5mC/5hmC on *HNF4A* P1 occurs, using *SFRS4* promoter as a control region. We could observe a constant level of 5hmC on *SFRS4* promoter in proliferative and differentiating cells (Figure 5B). *HNF4A P1* showed an increment of 5hmC at 1 week of differentiation as previously shown [Ancey et al., 2017]. A similar behaviour was found in *TCHP* and *RAB7A* regions (Figure 5B).

Next we asked how an adenosine derivative is able to disturb 5hmC levels triggering a differentiation delay on hepatocytes. To address this question we evaluated several components of the methionine cycle, which is the metabolic pathway responsible for biological *trans*-methylation reactions [Kharbanda, 2007]. Such reactions, including DNA methylation, could be modified by SAM availability as well as SAH levels [Auta et al., 2017; Garcea et al., 1989]. We previously showed that adenosine as well as IFC-305 are able to modulate SAM levels, favouring phospholipids methylation [Chagoya de Sanchez et al., 1991] and restoring global DNA methylation and 5hmC levels in a CCl_4_-mediated cirrhosis model [Rodriguez-Aguilera et al., 2018]. With this rationale, we evaluated the content of adenosine, SAM and SAH, through HPLC (Supplementary Figure S15). A trend to reduced SAM levels was observed in differentiating cells (Supplementary Figures S15A, S15B and S15E), consistent with global 5mC reduction at 1 week of HepaRG differentiation (Supplementary Figure S4A), with a non-significant trend to increase in SAH levels (Supplementary Figures S15A, S15B and S15F). Methylation index (SAM/SAH ratio) derived from these data, was lower and less variable in differentiating cells (Figure 5C). Higher and dispersed methylation index was observed in proliferative and IFC-305 exposed cells and correlates with high 5mC abundance genome wide (Supplementary Figure S4B). Adenosine measurement showed an increment in differentiating compared with proliferative cells, with the highest levels observed in differentiating + 1 mM IFC-305 cells, although not statistically significant (Supplementary Figure S15A, S15B, S15C and S15G; standard chromatogram is shown in Supplementary Figure S15D).

These results suggest that IFC-305 generates a methylation environment which competes with the demethylation wave associated with hepatocyte differentiation and highlights the importance of 5hmC enrichment in this process.

## Discussion

In this work, we have assessed the link between DNA methylation dynamics and the establishment of the hepatocyte-like transcription program after one week of HepaRG cells differentiation. We found that HepaRG cells at one week of differentiation have already triggered a hepatocyte transcriptional program, a time where there is a reduction of DNA methylation and 5hmC emerges genome-wide. While only 17% of DhMPs were associated to changes in gene expression, they were enriched in introns and depleted from promoters and CGIs. Finally, we found that perturbation of S-adenosylmethionine metabolism through an adenosine derivative, can reduce 5hmC enrichment associated to differentiation, triggering reduced levels of expression of hepatocyte markers.

Since its rediscovery in 2009 [Kriaucionis and Heintz, 2009; Tahiliani et al., 2009], 5hmC emerged as an intermediate of active DNA demethylation, but increasing evidence considers this modified cytosine as the “sixth base” because of its stability and its distinct functionality. Furthermore, 5hmC events correlate with key differentiation steps in mammal adult progenitors cells [Ecsedi et al., 2018]. Analysis of BS and OxBS from one week differentiating HepaRG cells as well as 5hmC immunostaining (Figures 2A and 2B, Supplementary Figure S2), showed a genome wide increase of this modified cytosine which corresponded with our previous findings on the *HNF4A P1* gene promoter [Ancey et al., 2017]. This behaviour of 5hmC enrichment has been described also in tissues from every germ layer, for example: enterocytes [Chapman et al., 2015; Kim et al., 2016], myocytes [Zhong et al., 2017], adipocytes [Dubois-Chevalier et al., 2014; Yoo et al., 2017] and neurons [Hahn et al., 2013; Li et al., 2017]. Hence, we could suggest that 5hmC enrichment in the last stage of cell differentiation is a mechanism widely present in human tissues; besides that, distribution of this mark at specific genomic locations could be characteristic of each cell type as it is observed in adult tissues [Lin et al., 2017].

It has been described that cell specification is accompanied by a stark transition in epigenetic landscape from a uniquely accessible state in pluripotency, to increasingly restrictive configurations in differentiated stage [Zhu et al., 2013]. However, 5hmC is often related to highly expressed cell specific genes [Li et al., 2016b; Lin et al., 2017]; hence in the present study, it is potentially playing a role in opening chromatin regions involved in hepatocyte specific transcription programs like metabolic pathways and with liver specialized functions as synthesis of complement components, regulation of lipids metabolism, platelet degranulation and oxidoreductase activity (Supplementary Figure S1A-S1D). Furthermore, we validated 5hmC enrichment on *HNF4A* and found DhMRs associated with *TET*s, and *IDH3G* (Supplementary Figures S5-S9), which could be related with a 5hmC feedback loop to reinforce differentiation. In line with these finding, it has been recently described that *TET1* is regulated through DNA methylation on its promoter [Li et al., 2016a]. This does not exclude that some other regulators could cooperate with this process or could also be generating restrictive chromatin context in non-hepatocyte related loci.

We found 5hmC enrichment in gene bodies (introns) and transcriptional features (Figure 3C and E) upon differentiation, features that partially agree with a report in adult mouse liver which indicates 5hmC is enriched in intragenic regions [Lin et al., 2017]. Similarly, in adult healthy liver, a specific 5hmC landscape has been described, which includes a bimodal 5hmC accumulation around TSS and an increased level along gene bodies, with another peak right after transcription termination [Li et al., 2016b]. Furthermore, we noticed that some DhMPs might be related with distal regulatory elements, as it could be the case for the 5hmC region associated with *TET3* (Supplementary Figure S9) which colocalizes with an enhancer for adult liver and overlaps with H3K4me1 and H3K27ac histone mark signals in HepG2 cells, as reported on the Epigenome Road Map Consortium and ENCODE, respectively.

In order to understand if acquisition of 5hmC is a key factor for differentiation, we assessed the effect of an adenosine derivative, IFC-305, which is able to modulate 5mC and 5hmC levels during experimental cirrhosis [Rodriguez-Aguilera et al., 2018]. We round that the hydroxymethylome of HepaRG cells exposed to IFC-305 presents an intermediate state between proliferative and differentiating cells (Figures 4 and 5, Supplementary Figure S13). This could be explained by the capacity of IFC-305 to increase SAM levels (Supplementary Figure S15E). SAM increase favours a methylating environment which could compete with DNA demethylation, therefore bringing a lower enrichment of 5hmC associated with attenuated differentiation. Contrary to our approach where cells are exposed to a SAM stimulating compound, a mouse model of methionine-choline-deficient diet which reduces SAM availability, did not cause a significant reduction of 5mC levels, instead inducing an up-regulation of Tet2, Tet3, thymine DNA glycosylase and apurinic/apyrimidic-endonuclease 1, and increased expression of Dnmt1 and Dnmt3a [Takumi et al., 2015]. Thus, while methylation dynamics could overcome SAM deficiency, SAM increase leads to reduced 5hmC enrichment with the consequent delay in differentiation evident at longer culture time (Supplementary Figure S13).

The possibility to modulate differentiation through regulation of 5hmC represents a relevant tool. In this sense, some studies indicate that one of the early alterations in HCC is a significant decrease of global genomic 5hmC, a behaviour also observed in hepatitis B virus infection [Liu et al., 2018], cirrhosis [Rodriguez-Aguilera et al., 2018], and during HSCs *trans*-differentiation to miofibroblast [Page et al., 2016]. Altered patterns of 5hmC are not only detectable in tumour-normal tissue pairs from resections or biopsies, but also in circulating DNA from cancer patients [Pfeifer and Szabo, 2018] suggesting that 5hmC is a potential biomarker for early cancer detection. Moreover, the possibility to modify cell identity through regulation of 5hmC distribution and levels represents an innovative pharmacological method to prevent or reverse certain pathologies.

## Conclusions

Our findings illustrate how differentiation of liver progenitor cells involves a demethylation process as well as a genome-wide increase in 5hmC. This 5hmC increase matches the enrichment of the same modified cytosine on promoter P1 of hepatocyte differentiation master regulator, *HNF4A*, and the activation of a liver transcriptional program at 1 week of differentiation. Moreover, a fraction of enriched 5hmC sites could be associated with over-expressed genes in differentiating cells. Furthermore, we found that cells exposed to an adenosine derivative presented lower 5hmC and did not reach levels of hepatocyte-expression markers observed in differentiating cells as well as presenting a less-differentiated phenotype. These results suggest a physiological role of 5hmC in acquisition of cell identity and support this modified cytosine as a potential biomarker for early detection of liver pathologies related with de-differentiation processes such as liver cancer.

## Methods

### Chemicals

Reagents were from Sigma Chemical Co. (St. Louis, MO). IFC-305 is the aspartate salt of adenosine: 2-aminosuccinic acid-2-(6-amino-9*H*-purin-9-yl)-5-(hydroxymethyl) tetrahydrofuran-3,4-diol (1:1). It was synthesized in the laboratory in agreement with UNAM patent 207422.

### HepaRG cell culture

Human HepaRG cells (Biopredic) were cultured as follows. Differentiating cells (4.4 × 10^4^/cm^2^) were grown for 1 week in William’s E medium (Gibco 12551-032) supplemented with 10% fetal bovine serum (Eurobio CVFSVF0001), 1x penicillin/streptomycin (Gibco 10378016), 5 μg/mL insulin (Sigma I9278) and 3 × 10^−5^ mM hydrocortisone (Sigma H0888) in 6 well plates (Figure 1), medium was replaced 24 h after seeding and once more after 3 days. For proliferative conditions, cells were seeded at low confluence, avoiding cell to cell contact to prevent cell differentiation. Treatment with IFC-305 began 24 h after cell seeding, and cells were re-treated at 3 day of culture when medium was replaced; this compound was directly solubilised in William’s E medium (Figure 1). Cells were harvested 7 days after seeding.

### Transcriptome analysis

RNA was isolated, directly from culture plate, with TRIzol (Invitrogen 15596018) and treated with DNase (Invitrogen 18068-015). Using the Illumina TotalPrep-96 RNA Amp Kit (Ambion 4393543), 500 ng of RNA was reverse transcribed into cDNA, which undergoes second strand synthesis to become a template for T7 RNA polymerase and thereby labelled with biotin-UTP. Hybridization of 2000 ng of labelled cDNA to the Illumina HumanHT-12 v3 (Illumina) were processed according to the manufacturer’s protocol. Slides were scanned immediately using Illumina BeadStation iScan (Illumina). Assays were performed in two independent cell cultures/condition.

### Quantitative RT-PCR

cDNA synthesis was performed from 2 μg of total RNA using M-MLV Reverse Transcriptase (Invitrogen 28025-013) and random primers. All quantitative PCR assays were performed independently in 3 cell cultures/condition in duplicate, with the delta-delta Ct method, using Mesa green qPCR 2x MasterMix Plus (Eurogentec 05-SY2X-06+WOU) on a CFX96 PCR system (Bio-Rad). Primers were described previously [Ancey et al., 2017].

### Cell survival evaluation

Differentiating and IFC-305 treated HepaRG cells cultured for 1 week were trypsinized and counted using TC20 automated cell counter (Bio-Rad) with dual chamber slides and trypan blue (Bio-Rad 1450003). Total cell and live cell numbers were determined automatically and % survival was determined by comparing live cells in each condition, with the mean of non-treated live cells which was considered as 100 %.

### Immunofluorescence

400 000 HepaRG cells/well were seeded in 6-wells plate with 4 coverslips/well, and were treated according to differentiation model with or without IFC-305. 100 000 cells were seeded in the same conditions during 24 h and were designated as proliferative cells. At different time points cells were washed with PBS, fixed in 4 % formaldehyde and washed twice with PBS. Primary antibodies for immunofluorescence were anti-HNF4α (Cell signalling 3113S), anti-5hmC (Active Motif 39769). After secondary antibodies, coverslips were washed and mounted on a slide with a mounting medium containing DAPI (Vectashield). Fluorescence was visualized with an Olympus Inverted Microscope model IX71 and images captured with Evolution/QImaging Digital Camera (Cybernetics). Data analysis was performed using ImageJ (National Institutes of Heatlh).

### Oxidative Bisulfite and Methylation Bead Arrays

For genomic DNA isolation, 500 μL of lysis buffer (50 Mm Tris-HCl pH 8, 100 mM EDTA, 100 mM NaCl, 1% SDS and 0.5 mg/mL proteinase K) were added to wells from 6-well plates, previously washed twice with PBS 1x. Cells were detached by pipetting and suspension was incubated 2 h at 55°C. 200 μL of 6 M NaCl was added, sample was mixed and centrifuged 10 min at full speed. Supernatant was recovered and DNA was precipitated with 500 μL isopropanol, washed with 500 μL 70% ethanol and dried by inverting the tube on a tissue. DNA was resuspended with 25 μL injectable water. 1000 ng DNA was oxidative bisulfite- and conventional bisulfite-converted using TrueMethyl Seq Kit (Cambridge Epigenetix) according to manufacturer’s instructions. This technique oxidises 5hmC to 5fC which is read as thymine (non-modified cytosine and 5mC are read as thymine and cytosine respectively, as in conventional bisulfite) [Booth et al., 2013]. Converted DNA was analysed with MethylationEPIC arrays (Illumina) using recommended protocols for amplification, labelling, hybridization, and scanning. For proliferative and differentiating cells, each methylation analysis was performed in two independent culture wells and for IFC-305 treatment in triplicates.

### Bioinformatic Analyses

Raw expression and methylation data were imported and processed using R/Bioconductor packages for Illumina bead arrays [Du et al., 2008]. Data quality was inspected using boxplots for the distribution of expression signals, and inter-sample relationship using multidimensional scaling plots and unsupervised clustering. For methylation data, we removed low quality probes with a detection P value > 0.01 in more than 10% of the samples. Following swan (Subset-quantile Within Array) normalization [Maksimovic et al., 2012] implemented in the minfi package [Aryee et al., 2014], 5hmC “peaks” were identified by subtracting the signal of oxidative bisulfite from the signal of conventional bisulfite. Next, we performed Maximum Likelihood Estimate (MLE) of 5mC and 5hmC (oxBS.MLE) with the ENmix package [Xu et al., 2016]. Using a binomial model at each CpG locus in each sample, oxBS.MLE outputs a matrix of MLEs of 5mC levels and a matrix of MLEs of 5hmC levels, setting as NA any negative value. To define differentially expressed genes (DEGs), differentially methylated positions (DMPs) and differentially hydroxymethylated positions (DhMPs), we modelled experimental conditions as categorical variables in a linear regression using an empirical Bayesian approach [Smyth, 2004]. Differentially methylated and hydroxymethylated regions (DMRs and DhMRs, respectively) were identified with the DMRcate package using the recommended proximity-based criteria [Peters et al., 2015]. Comparisons with an FDR-adjusted P value below 0.05 were considered statistically significant. In addition, only DhMPs with at least 10% change in 5hmC were considered as significant for any given comparison, as this has been suggested as a threshold for the sensitivity of this technique [Skvortsova et al., 2017]. DEGs, DMPs, and DMRs were further analyzed to determine functional pathways and ontology enrichment using EnrichR [Chen et al., 2013]. Genomic context annotations, including chromatin states (ChromHMM) were performed with the ChIPseeker [Yu et al., 2015] and Annotatr [Cavalcante and Sartor, 2017] packages. EnrichedHeatmap package [Gu et al., 2018] was used for summary heatmaps of genomic context. All expression and methylation data will be deposited to the Gene Expression Omnibus repository.

### DNA immunoprecipitation

80 uL of aqueous genomic DNA (100 ng/uL) were sonicated in Covaris microTUBE AFA Fiber Pre-Slit Snap-Cap 6×16mm with Covaris sonicator S220 to obtain fragments between 400-800 bp (Temperature < 7°C, peak power 105.0, duty factor 5.0, Cycles/Burst 200 and time 40 s). 130 ng DNA was used per immunoprecipitation. Auto hMeDIP kit (Diagenode C02010033) was used according to manufacturer instructions. Precipitation was adjusted to 15 h for mixing at 4°C and middle mix speed; for washes mixing time was 8 min, at 4°C and middle mix speed. Immunoprecipitations were performed independently in 3 cell cultures/condition. 5 uL of immunoprecipitated hydroxymethyl DNA was analysed for enriched 5hmC regions according to oxBS data (regions with at least 3 CpGs containing a DhMP with 10 % difference between proliferative and differentiating cells (Delta 10%)), *SFRS4* gene promoter was used as control of non-5hmC enrichment. All quantitative PCR assays were performed in duplicate with Mesa green qPCR 2x MasterMix Plus (Eurogentec 05-SY2X-06+WOU) on a CFX96 PCR system (Bio-Rad) using primers indicated in Supplementary Table S1. 5hmC enrichment was determined as % (hmeDNA-IP/ Total input) as follows: % (hmeDNA-IP/ Total input) = 2^[(Ct(10%input) - 3.32) - Ct(hmeDNA-IP)]x 100%, where 2 is the amplification efficiency; Ct (hmeDNA-IP) and Ct (10%input) are threshold values obtained from exponential phase of qPCR for the hydroxymethyl DNA sample and input sample respectively; the compensatory factor (3.32) is used to take into account the dilution 1:10 of the input.

### Adenosine, S-adenosylmethionine and S-adenosylhomocysteine quantification by HPLC

Adenosine, SAM and SAH levels were determined modifying protocols previously described [Hernandez-Munoz et al., 1984; Korinek et al., 2013]. Briefly, 10*10^6^ proliferating HepaRG cells were seeded in a 15 cm Petri dish; 24h after seeding, cells were either cultured with or without IFC-305 for 1 week to get differentiate, or collected for the Proliferative condition. Next, cells were frozen with liquid nitrogen and preserved at −70°C. Frozen cells were harvested with 1.5 mL 0.03% trifluoroacetic acid in 90% MeOH. Samples were incubated 10 min at room temperature and passed through a Dounce homogenizer. Obtained solutions were sonicated twice for 1 min in a Bransonic 220 and centrifuged at 16 000 g for 13 min at 4°C. Supernatants were recovered and centrifuged until dry in a speedvac at 45°C. Samples were reconstituted with 200 μL MilliQ water and protein concentration was determined by Bradford assay. Samples were diluted to 1.2 ug/uL protein and 100 μL 1.6 M HClO_4_ was added to 300 μL of diluted sample (0.4 M HClO_4_ final concentration) before incubating for 10 min on ice. 10 μL of 5 M K_2_CO_3_ was added to samples and incubated for 10 min on ice. Sample dilutions were centrifuged at 14 000 rpm during 5 min in a Eppendorf 5415C Micro-Centrifuge. Recovered supernatants were filtered using Phenex PTFE 4 mm syringe filters and 95 μL supernatant + 5 μL of 400 mM adenosine and 400 mM SAM standards mix were injected to HPLC Knauer E4310 (SAM standard contains SAH contamination, which was determined using SAH standard curve (data not shown)). Samples were separated in an ACE 5 C 18 column (150 × 4.6 mm) (Advanced Chromatpgraphy Technologies LTD) using a mobile phase consisting of 8 mM sodium heptanesulfonate, 40 mM ammonium phosphate monobasic and 15 % methanol pH 3 without gradient, at 1 mL/min flux, detection was measured at 254 nm absorbance, and separation total runtime of 60 min. Peaks were analyzed using EUROCHROME for windows version 3.05 (Knauer GmbH).

## Abbreviations

5mC: 5-methylchytosine
5hmC: 5-hydroxymethylcytosine
BS: bisulfite sequencing
CGI: CpG island
DMP: differentially methylated position
DMR: differentially methylated region
DNMTs: DNA methyl-transferases
HCC: Hepatocellular carcinoma
iCCA: intrahepatic cholangiocarcinoma
oxBS: oxidative bisulfite sequencing
SAM: s-adenosylmethionine
SAH: S-adenosylhomocysteine
TETs: Tet Methylcytosine Dioxigenases
TF: transcription factor
TSS: transcription start sites

## Availability of data and material

Datasets generated during the current study will be uploaded to the GEO repository.

## Ethics approval and consent to participate

Not applicable.

## Consent for publication

Not applicable.

## Authors’ contributions

JRRA carried out the experiments and co-wrote the first draft of the manuscript; SE performed sample preparations for HT-12 array and advice in MeDIP assays; MPC performed immunofluorescence assays and provided technical assistance in the progenitor differentiation experiments and qPCR assays; CG performed additional validations and co-wrote the first draft of the manuscript; MDL standardized and provided technical assistance in HPLC experiments; NGC and RPCV performed additional differentiation assays and contributed to HPLC experiments; IC and FRT contributed to discussion of results and provided conceptual assistance; VCS conceived the study, provided conceptual assistance and supervised experiments; HHV conceived the study, carried out statistical and bioinformatics analysis, supervised experiments, coordinated the project and wrote the manuscript. All authors discussed the results and manuscript text.

## Acknowledgements

The authors thank Cyrille Cuenin (Epigenetics Group, IARC, Lyon) and Severine Croze (Claude Bernard, University Lyon 1) for their help with the HT-12 array and MethylEPIC assays. The authors also thank Florence Le Calvez-Kelm and Geoffroy Durand (Genetic Cancer Susceptibility Group, IARC, Lyon) for Covaris Sonicator supply, Zdenko Herceg (Epigenetics Group, IARC, Lyon) for facilitating the infrastructure to realize hMeDIP experiments, Ramón González García-Conde (Research Center on Cellular Dynamics, UAEM, Mexico) for HepaRG cell line donation for preliminary assays, Emilio-Rojas del Castillo (Department of Genomic Medicine and Environmental Toxicology, IIB-UNAM, Mexico) for his advice during JRRA PhD studies, Pedro Valencia Mayoral (Planning Direction, Hospital Infantil de México “Federico Gómez”) for training JRRA on liver histopathology during PhD Studies, Athena Sklias and Andrea Halaburkova (Epigenetics Group, IARC, Lyon) for their enriching discussions during laboratory work, and Aurelie Salle (Epigenetics Group, IARC, Lyon), Georgina Guerrero-Avendaño and Rosario Pérez-Molina (Department of Molecular Genetics, IFC-UNAM, Mexico) for their technical assistance.

