## Supplementary Data for "Genome-wide 5-hydroxymethylcytosine (5hmC) emerges at early stage of *in vitro* hepatocyte differentiation"

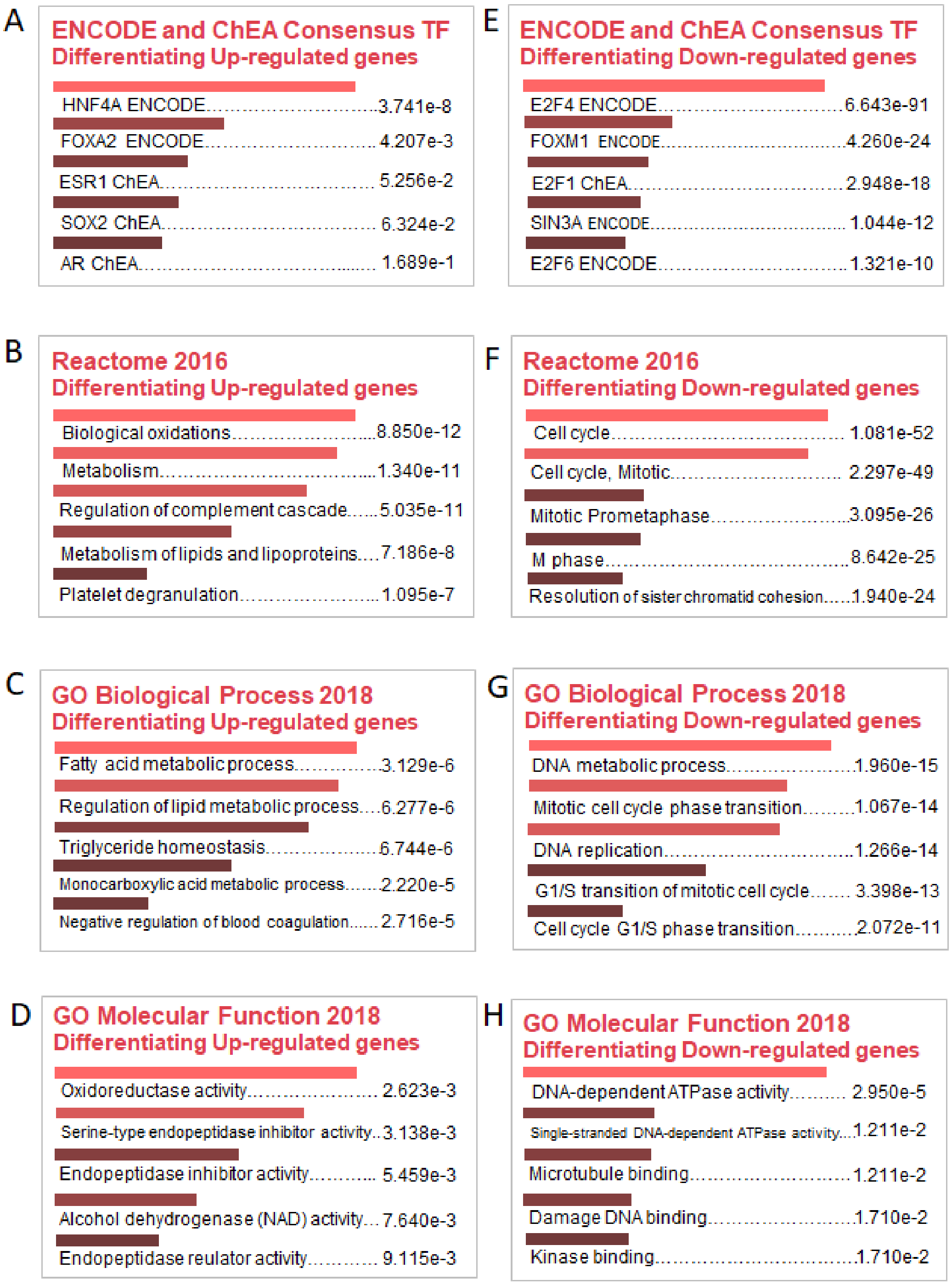


**Supplementary Figure S1. Gene ontologies from differentiating cells.** (A and E) Consensus transcription factors (TF) from Encyclopedia of DNA elements (ENCODE) and ChIP-X Enrichment Analysis (ChEA) of differentiating up-regulated genes and down regulated genes respectively. (B-D) Ontologies related with up-regulated genes from differentiating cells. (F-H) Ontologies related with down-regulated genes from differentiating cells. Data by EnrichR, number showed tissue-associated adjusted p-value.

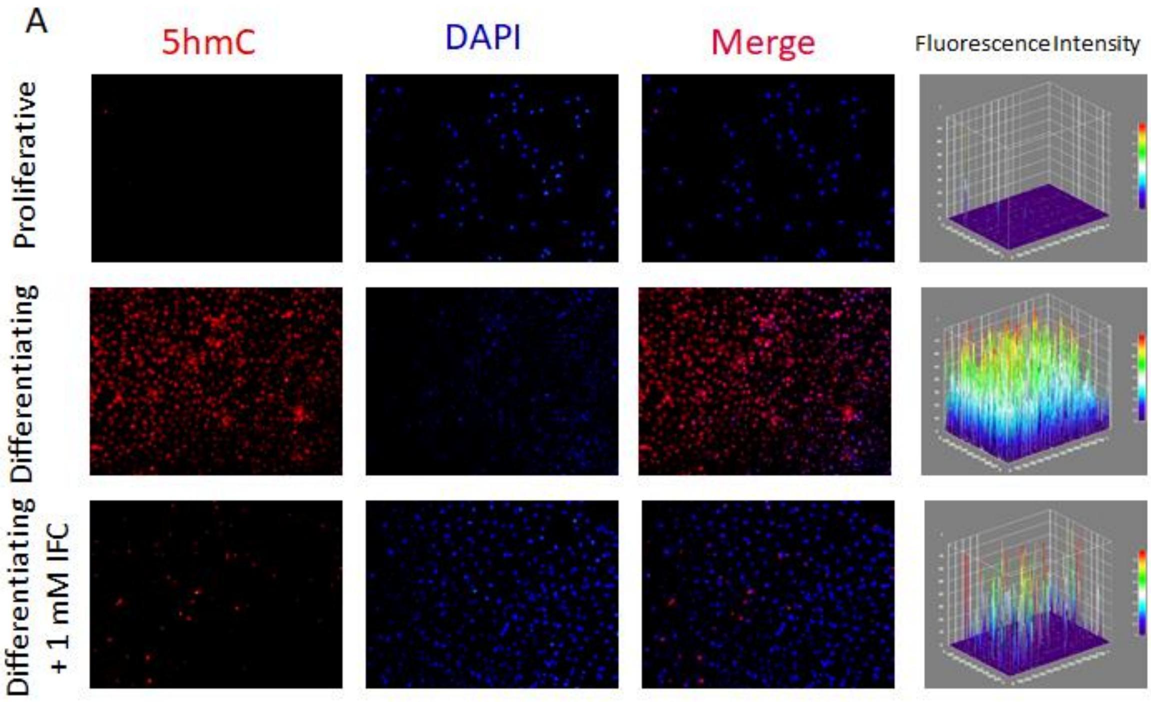


**Supplementary Figure S2. 5hmC appears in differentiating cells.** Immunofluorescence of 5hmC in proliferative (top panel), differentiating (middle panel) and differentiating + IFC-305 (bottom panel). Representative images from 3 fields/condition are shown, as well as measurement of fluorescence intensity.

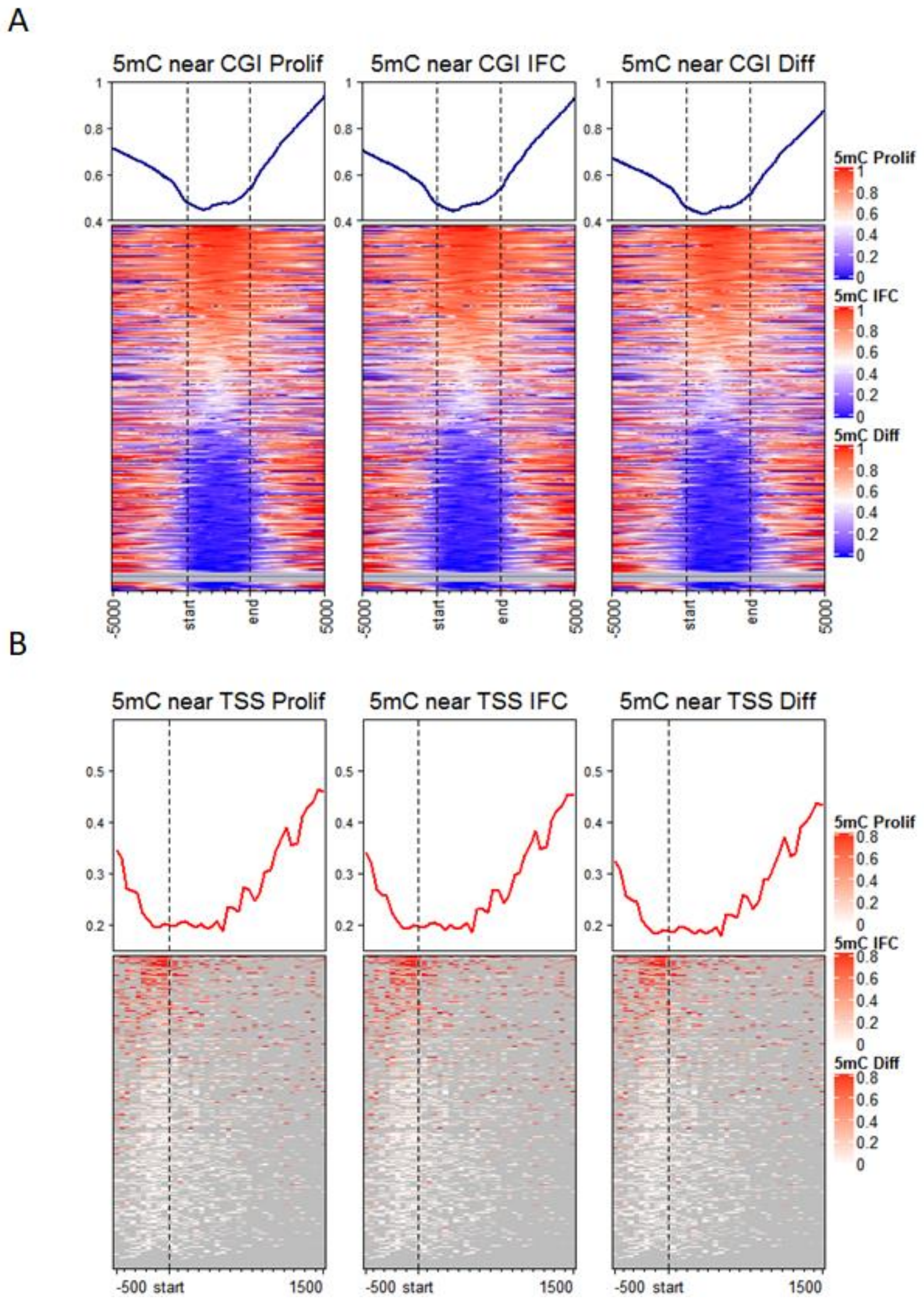


**Supplementary Figure S3. 5mC distribution on CpG islands and TSS during hepatocyte differentiation.** Global distribution of 5hmC for one representative sample of each condition, according to CpG islands (A) and transcription start sites (B). In both cases, 5hmC levels are averaged across all hg19-annotated genomic regions. Two independent cultures were used for proliferative and differentiating cells.

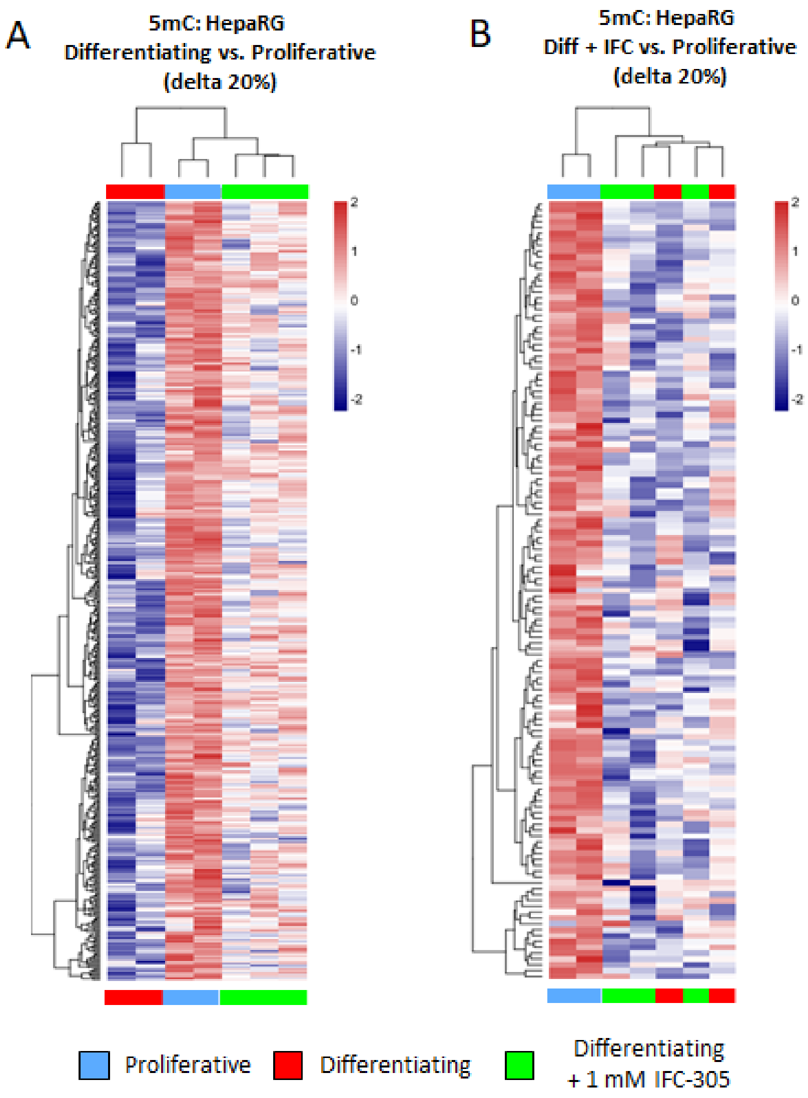


**Supplementary Figure S4. Differentiating process involves a genome-wide demethylation.** (A) Heatmap showing methylome comparison between differentiating and proliferative cells. (B) Heatmap showing methylome comparison between differentiating + IFC-305 and proliferative cells. Differentially methylated positions (DMPs) were filtered by the magnitude of change in methylation (delta beta) of at least 20% and p-adjusted value $< 0.05$. Two independent cultures were used for proliferative and differentiating cells, and three independent cultures for differentiating + 1 mM IFC-305.

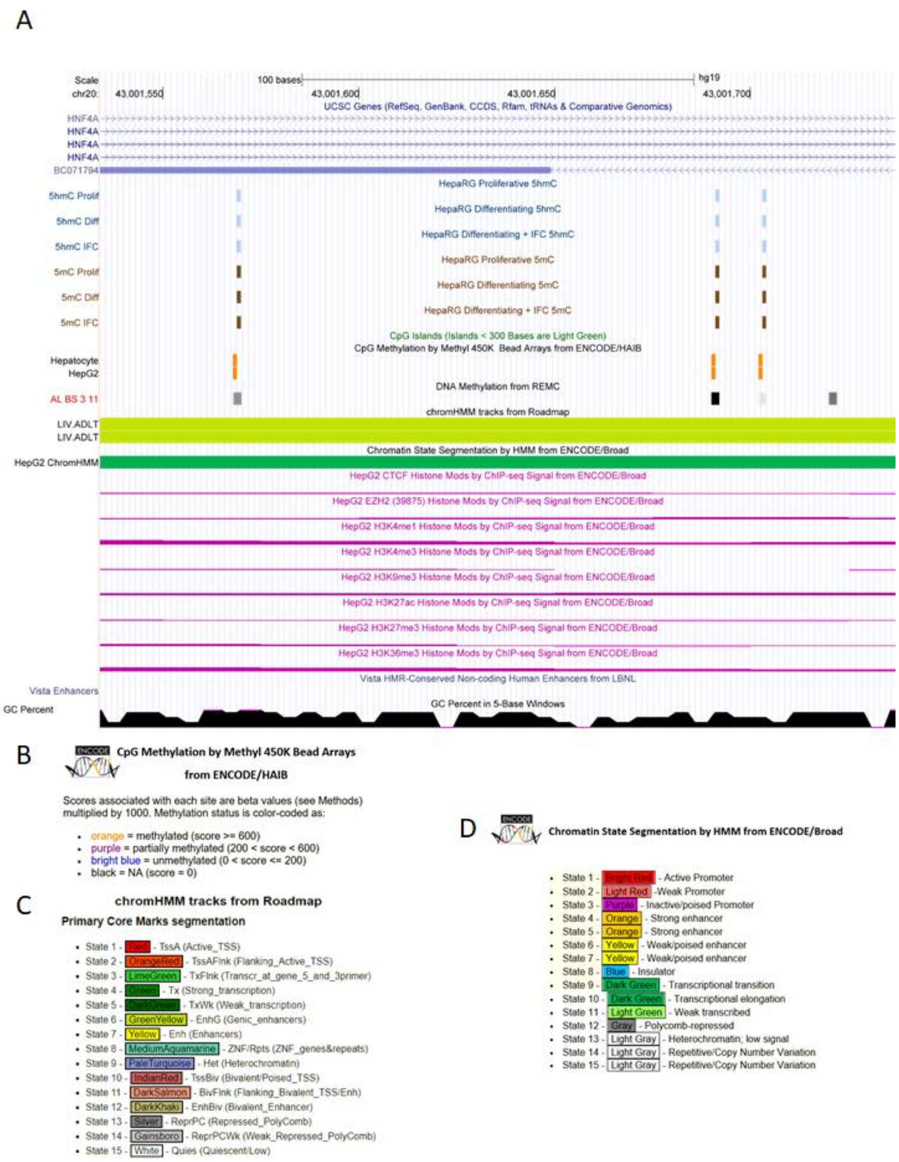


**Supplementary Figure S5. 5hmC region on *HNF4A*.** UCSC Genome Browser screen, presenting region chr20:43001569-43001703, which flank 5hmC-enriched 3 CpG sites on *HNF4A* gene. 5hmC data is shown in blue and 5mC data in brown; color intensity is proportional to enrichment degree. Proliferative (Prolif), Differentiating (Diff), Differentiating + IFC (IFC). Some additional tracks are shown: CpG islands < 300 bp; CpG methylation by Methyl 450K Arrays from ENCODE/HAIB of Hepatocytes and HepG2 cell line, color-code is denoted in (B); DNA Methylation from Roadmap Epigenomics Mapping Consortium (REMC) of adult liver bisulfite sequencing (AL BS), gray intensity is proportional to enrichment degree; Chromatin State Segmentation by Hidden Markov Model (ChromHMM) of adult liver (LIV ADLT) and HepG2 cell line, color-code is denoted in (C and D) respectively; HepG2 tracks for architectural protein CTCF, EZH2, and histone marks as H3K4me1 (enhancers), H3K4me3 (promoters), H3K36me3 (elongation), H3K9me3 and H3K27me3 (close chromatin marks), H3K27ac (open chromatin mark) from ENCODE/Broad; Vista Enhancer Handbook and Methods (HMR) at the Lawrence Berkeley National Laboratory (LBNL), peaks denote enrichment; and CG percent, peaks indicates abundance.

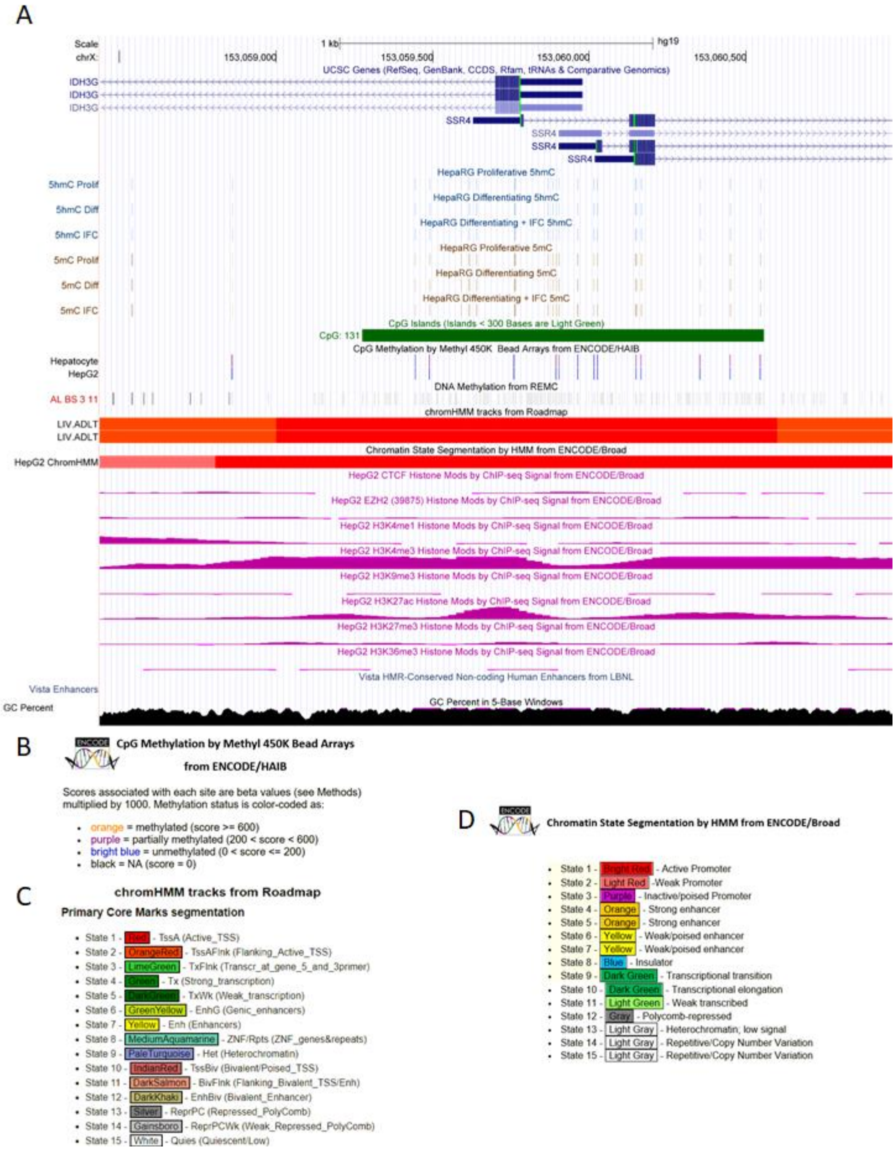


**Supplementary Figure S6. 5hmC region on *IDH3G*.** UCSC Genome Browser screen, presenting region chrX:153058860-153060546, which flank 5hmC-enriched 21 CpG sites on *IDH3G* gene. 5hmC data is shown in blue and 5mC data in brown; color intensity is proportional to enrichment degree. Proliferative (Prolif), Differentiating (Diff), Differentiating + IFC (IFC). Some additional tracks are shown: CpG islands < 300 bp; CpG methylation by Methyl 450K Arrays from ENCODE/HAIB of Hepatocytes and HepG2 cell line, color-code is denoted in (B); DNA Methylation from Roadmap Epigenomics Mapping Consortium (REMC) of adult liver bisulfite sequencing (AL BS), gray intensity is proportional to enrichment degree; Chromatin State Segmentation by H*idden Markov* Model (ChromHMM) of adult liver (LIV ADLT) and HepG2 cell line, color-code is denoted in (C and D) respectively; HepG2 tracks for architectural protein CTCF, EZH2, and histone marks as H3K4me1 (enhancers), H3K4me3 (promoters), H3K36me3 (elongation), H3K9me3 and H3K27me3 (close chromatin marks), H3K27ac (open chromatin mark) from ENCODE/Broad; Vista Enhancer Handbook and Methods (HMR) at the Lawrence Berkeley National Laboratory (LBNL), peaks denote enrichment; and CG percent, peaks indicates abundance.

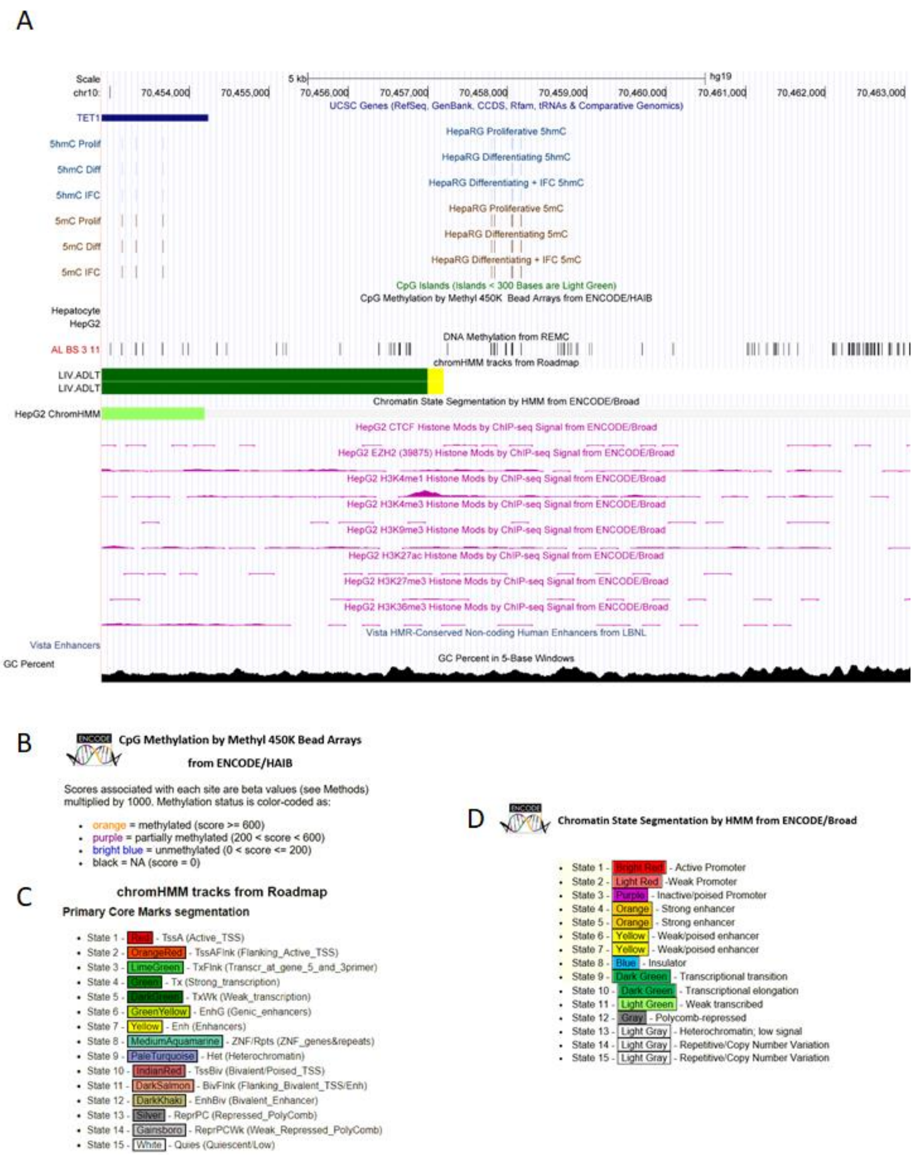


**Supplementary Figure S7. 5hmC region on *TET1.*** UCSC Genome Browser screen, presenting region chr10:70457799-70458175, which flank 5hmC-enriched 5 CpG sites on *TET1* gene. 5hmC data is shown in blue and 5mC data in brown; color intensity is proportional to enrichment degree. Proliferative (Prolif), Differentiating (Diff), Differentiating + IFC (IFC). Some additional tracks are shown: CpG islands < 300 bp; CpG methylation by Methyl 450K Arrays from ENCODE/HAIB of Hepatocytes and HepG2 cell line, color-code is denoted in (B); DNA Methylation from Roadmap Epigenomics Mapping Consortium (REMC) of adult liver bisulfite sequencing (AL BS), gray intensity is proportional to enrichment degree; Chromatin State Segmentation by H*idden Markov* Model (ChromHMM) of adult liver (LIV ADLT) and HepG2 cell line, color-code is denoted in (C and D) respectively; HepG2 tracks for architectural protein CTCF, EZH2, and histone marks as H3K4me1 (enhancers), H3K4me3 (promoters), H3K36me3 (elongation), H3K9me3 and H3K27me3 (close chromatin marks), H3K27ac (open chromatin mark) from ENCODE/Broad; Vista Enhancer Handbook and Methods (HMR) at the Lawrence Berkeley National Laboratory (LBNL), peaks denote enrichment; and CG percent, peaks indicates abundance.

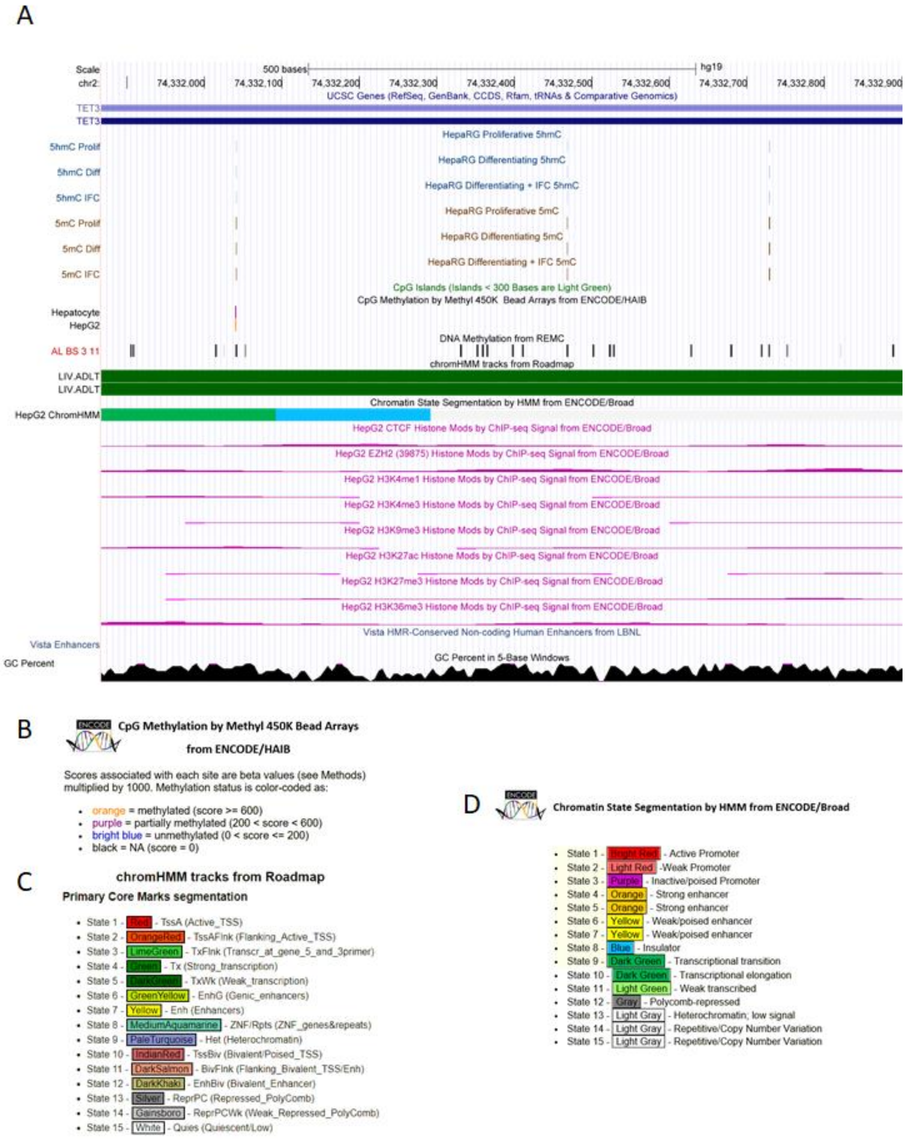


**Supplementary Figure S8. First 5hmC region on *TET3*.** UCSC Genome Browser screen, presenting region chr2:74332041-74332729, which flank 5hmC-enriched 3 CpG sites on *TET3* gene. 5hmC data is shown in blue and 5mC data in brown; color intensity is proportional to enrichment degree. Proliferative (Prolif), Differentiating (Diff), Differentiating + IFC (IFC). Some additional tracks are shown: CpG islands < 300 bp; CpG methylation by Methyl 450K Arrays from ENCODE/HAIB of Hepatocytes and HepG2 cell line, color-code is denoted in (B); DNA Methylation from Roadmap Epigenomics Mapping Consortium (REMC) of adult liver bisulfite sequencing (AL BS), gray intensity is proportional to enrichment degree; Chromatin State Segmentation by H*idden Markov* Model (ChromHMM) of adult liver (LIV ADLT) and HepG2 cell line, color-code is denoted in (C and D) respectively; HepG2 tracks for architectural protein CTCF, EZH2, and histone marks as H3K4me1 (enhancers), H3K4me3 (promoters), H3K36me3 (elongation), H3K9me3 and H3K27me3 (close chromatin marks), H3K27ac (open chromatin mark) from ENCODE/Broad; Vista Enhancer Handbook and Methods (HMR) at the Lawrence Berkeley National Laboratory (LBNL), peaks denote enrichment; and CG percent, peaks indicates abundance.

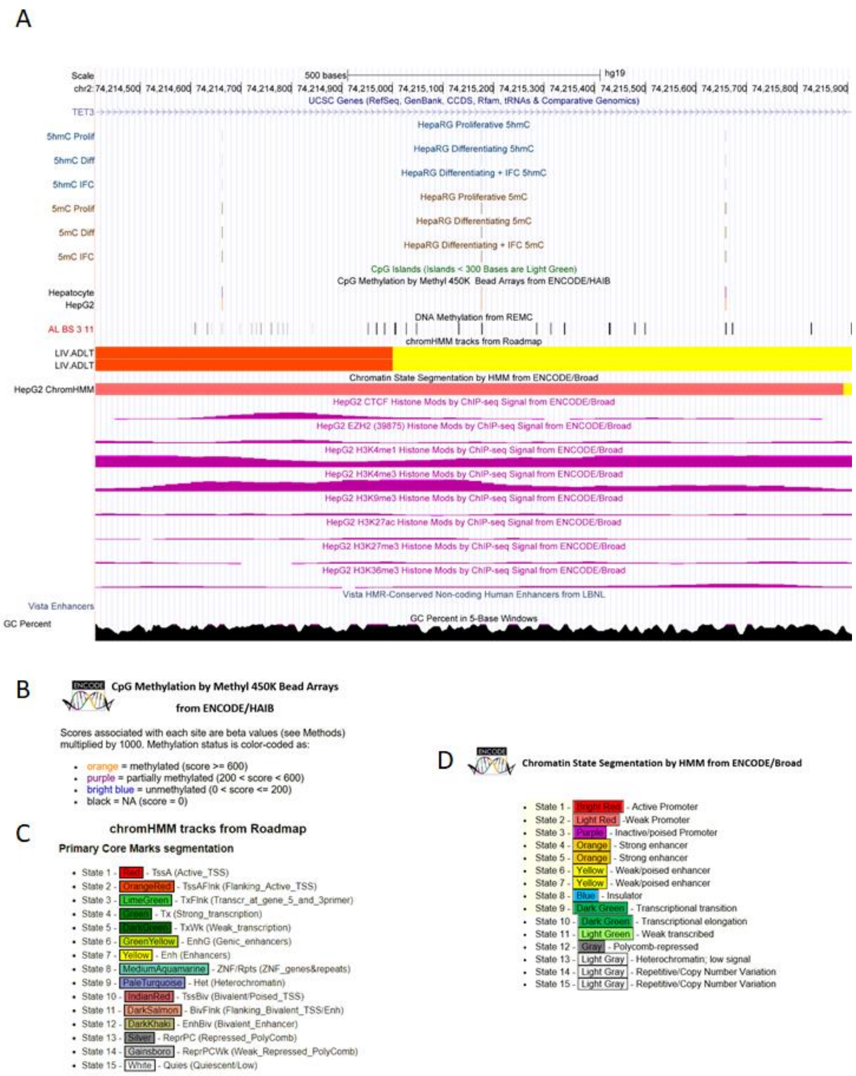


**Supplementary Figure S9. Second 5hmC region on *TET3*.** UCSC Genome Browser screen, presenting region chr2:74214662-74215659, which flank 5hmC-enriched 3 CpG sites on *TET3* gene. 5hmC data is shown in blue and 5mC data in brown; color intensity is proportional to enrichment degree. Proliferative (Prolif), Differentiating (Diff), Differentiating + IFC (IFC). Some additional tracks are shown: CpG islands < 300 bp; CpG methylation by Methyl 450K Arrays from ENCODE/HAIB of Hepatocytes and HepG2 cell line, color-code is denoted in (B); DNA Methylation from Roadmap Epigenomics Mapping Consortium (REMC) of adult liver bisulfite sequencing (AL BS), gray intensity is proportional to enrichment degree; Chromatin State Segmentation by *Hidden Markov* Model (ChromHMM) of adult liver (LIV ADLT) and HepG2 cell line, color-code is denoted in (C and D) respectively; HepG2 tracks for architectural protein CTCF, EZH2, and histone marks as H3K4me1 (enhancers), H3K4me3 (promoters), H3K36me3 (elongation), H3K9me3 and H3K27me3 (close chromatin marks), H3K27ac (open chromatin mark) from ENCODE/Broad; Vista Enhancer Handbook and Methods (HMR) at the Lawrence Berkeley National Laboratory (LBNL), peaks denote enrichment; and CG percent, peaks indicates abundance.

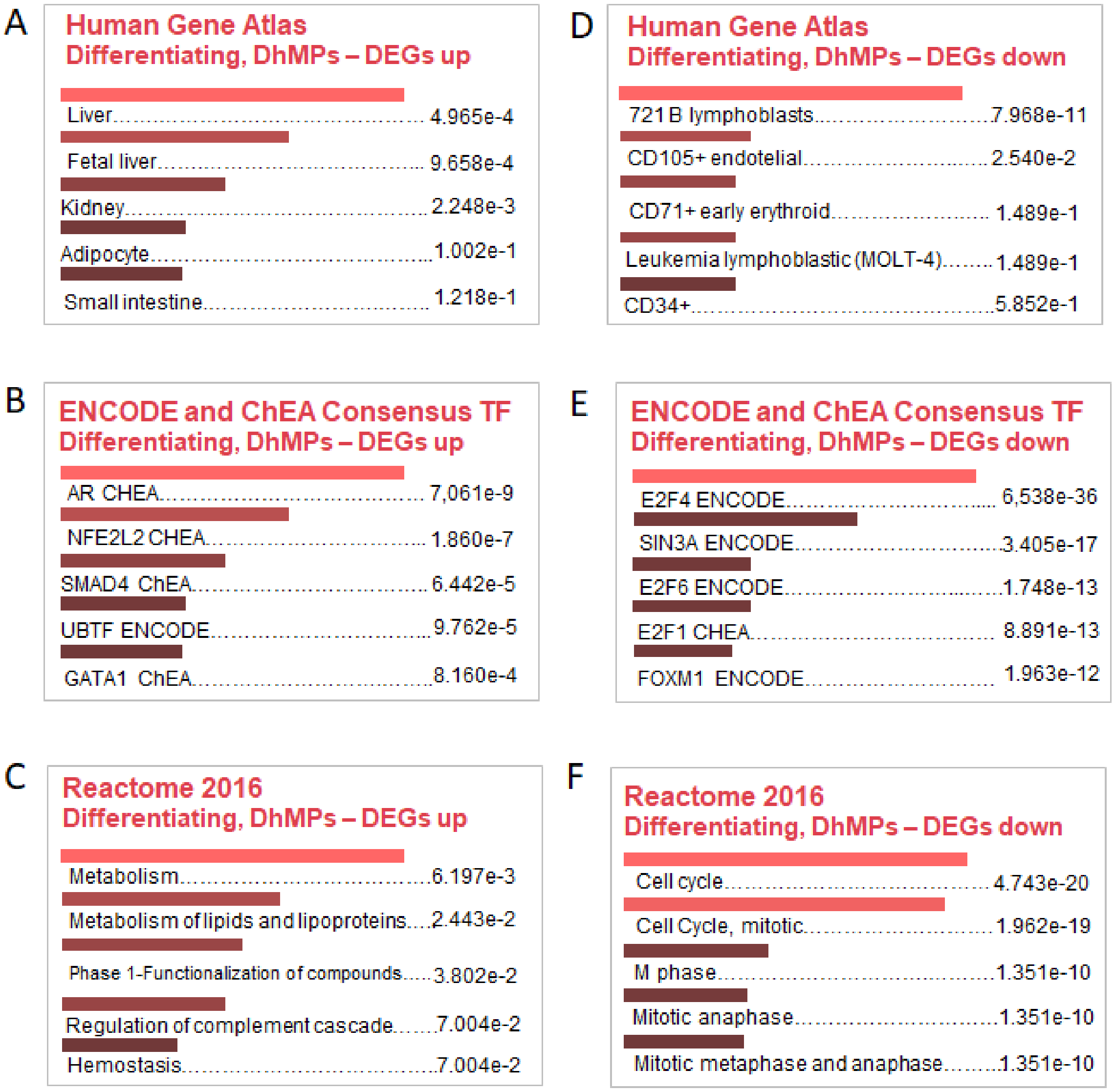


**Supplementary Figure S10. Ontologies from overlaps between DhMPs and DEGs.** (A) Cells/tissues associated with differentiating enriched 5hmC positions and up-regulated genes comparing Differentiating vs. Proliferative. (B) Consensus transcription factors (TF) from ENCODE ChEA of differentiating enriched 5hmC positions and up-regulated genes. (C) Ontologies related with differentiating enriched 5hmC positions and up-regulated genes. (D) Cells/tissues associated with differentiating enriched 5hmC positions and down-regulated genes comparing Differentiating vs. Proliferative. (E) Consensus transcription factors (TF) from ENCODE ChEA of differentiating enriched 5hmC positions and down-regulated genes. (F) Ontologies related with differentiating enriched 5hmC positions and down-regulated genes. Data from EnrichR, number showed tissue-associated adjusted p-value.

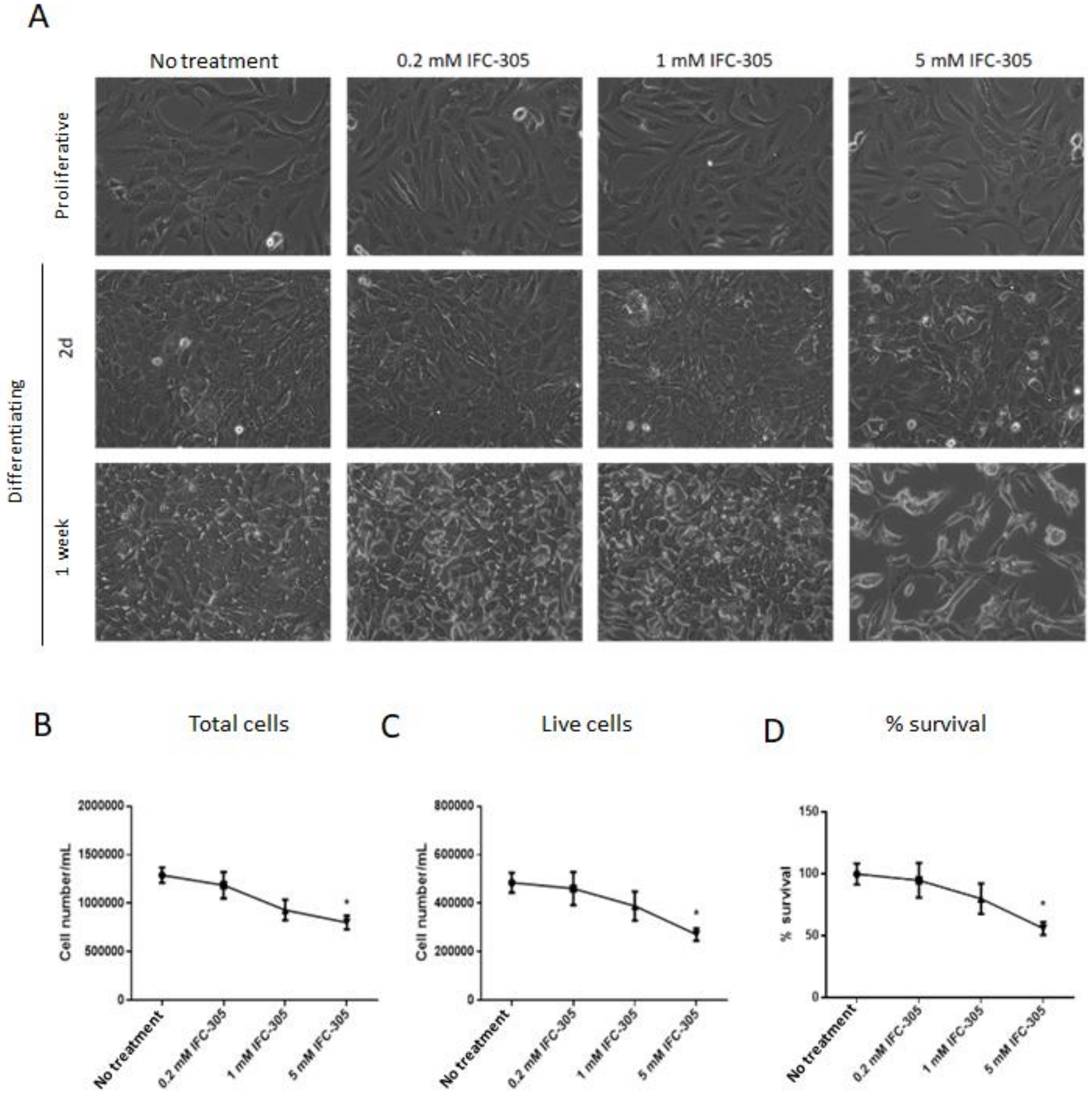


**Supplementary Figure S11. Low concentrated adenosine derivative did not alter HepaRG cell viability.** (A) HepaRG cell phenotype along 1 week of differentiation. 20x representative images of each group are showed. It is possible to observe 5 mM IFC-305 treatment retains elongated proliferative phenotype and 0.2 and 1 mM IFC-305 did not modify cell phenotype compared with differentiating cells. (B) Total cell number. (C) Viable cell number. (D) Survival percentage. Data represent mean ± SEM of seven independent cultures. *Statistical difference ($p < 0.05$) compared with differentiating cells.

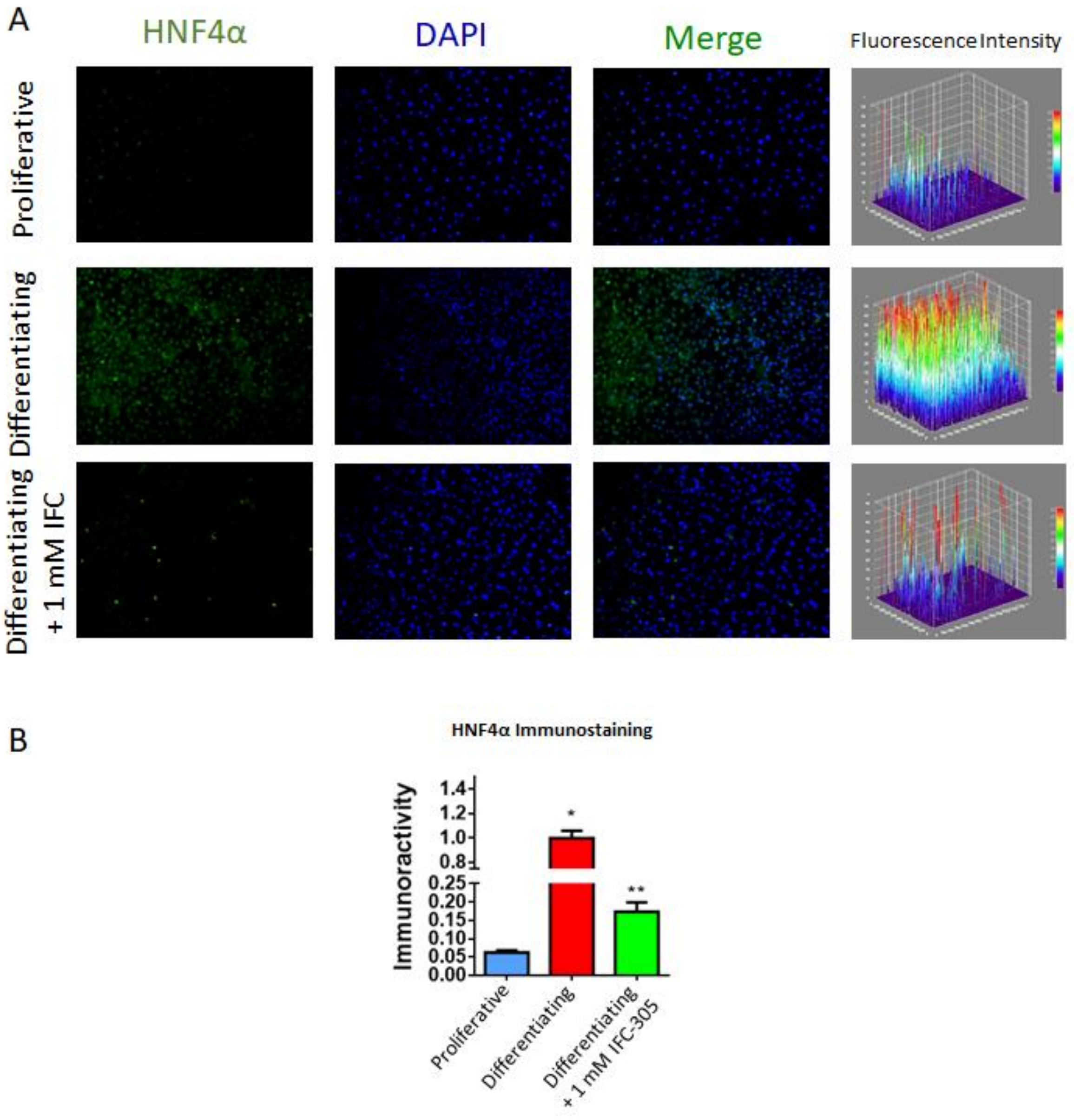


**Supplementary Figure S12. HNF4α increases in differentiating cells.** (A) Immunofluorescence of HNF4α in proliferative (top panel), differentiating (middle panel) and differentiating + IFC-305 (bottom panel). Representative images from 3 fields/condition are shown. (B) Quantification of immunofluorescence HNF4α positive signal. Data represent mean ± SEM from 3 fields/group. *Statistical difference ($p < 0.05$) when compared with proliferative cells. **Statistical difference ($p < 0.05$) compared with differentiating cells.

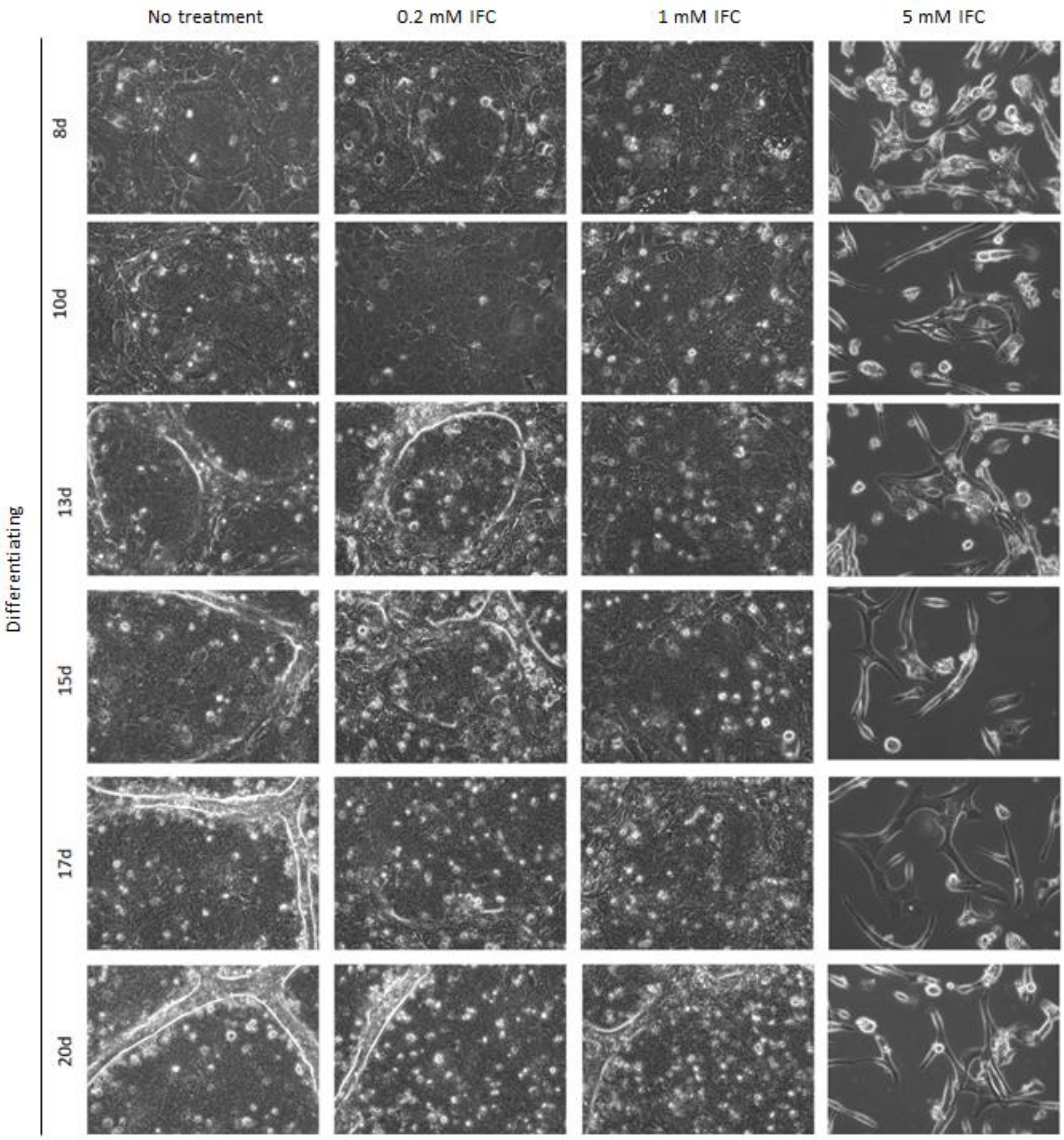


**Supplementary Figure S13. 1mM IFC-305 differentiating exposed cells presents a less differentiated phenotype.** 20 days time-lapse images showing HepaRG differentiating phenotype with an increasing gradient of IFC-305. Representative images from seven independent cultures. Representative 20x magnification images from seven independent cultures are shown.

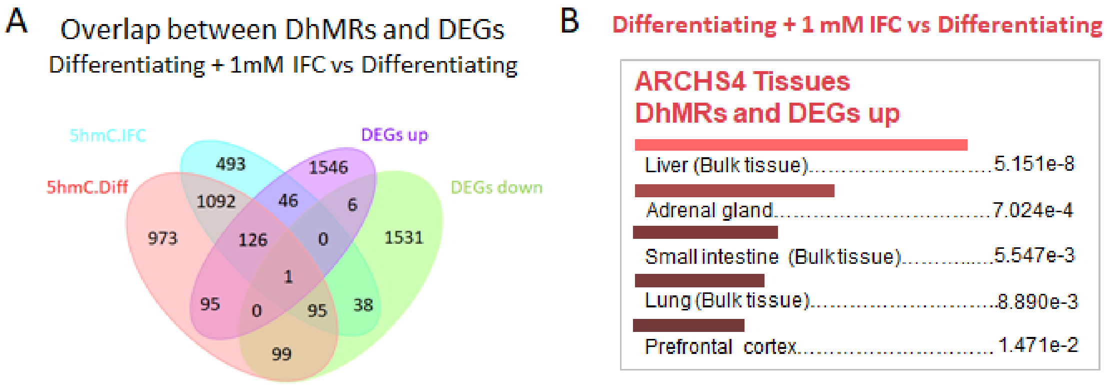


**Supplementary Figure S14. Some differential hydroxymethylated regions in IFC-305 treated differentiating cells, were related with differential overexpressed genes.** (A) Comparison between 5hmC differentiated regions (DhMRs) in IFC-305 exposed cells and DEGs in differentiating cells. (B) Ontologies related with differentiating + IFC-305 enriched 5hmC regions and up-regulated genes in differentiating cells. Data from EnrichR, number showed tissue-associated adjusted p-value.

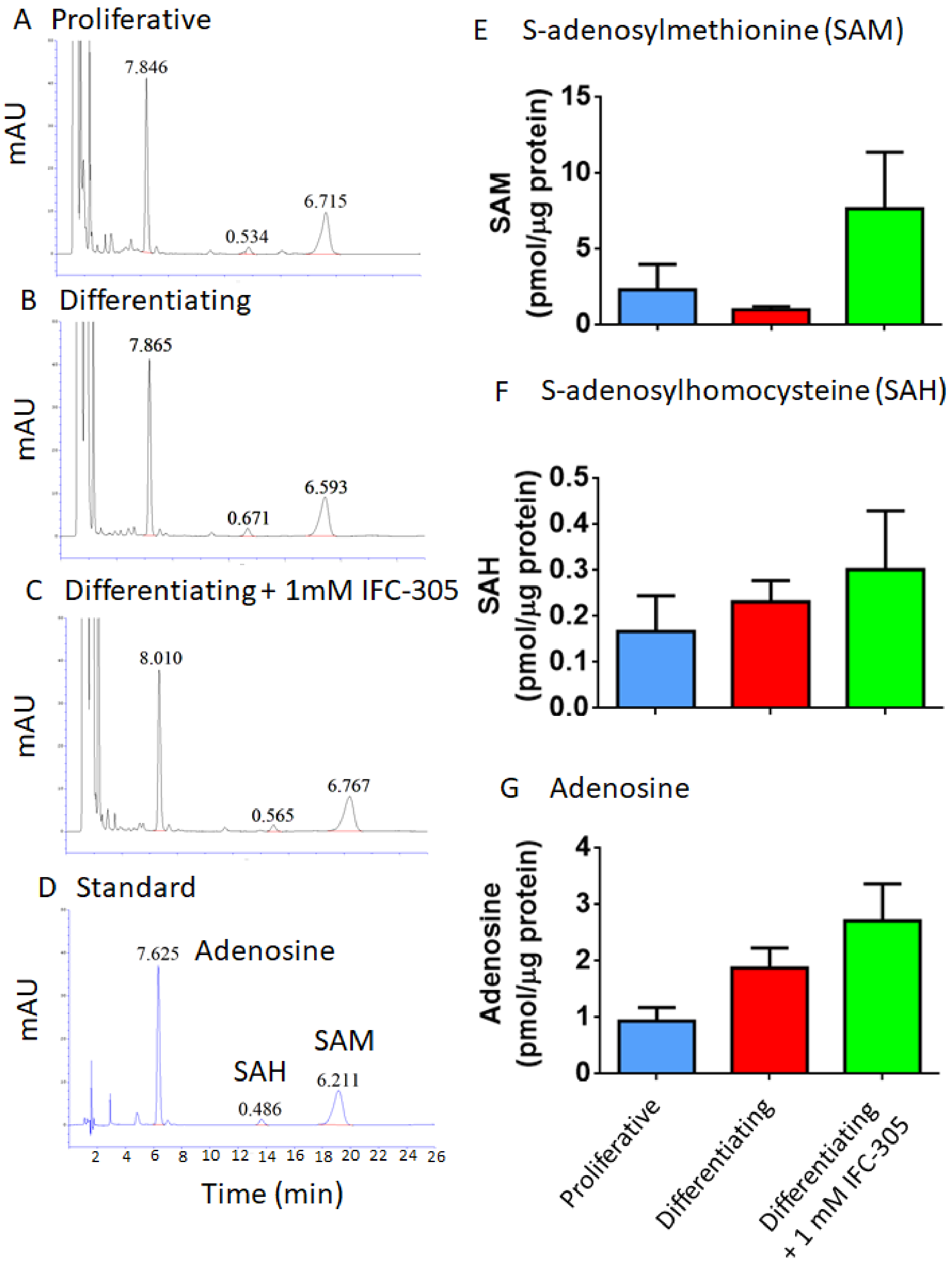


**Supplementary Figure S15. S-adenosylmethionine trends to decrease with HepaRG differentiation.** (A-D) Representative chromatograms identifying adenosine, S-adenosylhomocysteine (SAH) and S-adenosylmethionine (SAM) through HPLC in analyzed conditions and generated by standards, numbers indicate area under curve, first 26 min separation runtime are shown. (E-G) Quantification of each analyte. Data represent mean ± SEM from 4 cultures/condition.
